# Host phylogeny and host ecology structure the mammalian gut microbiota at different taxonomic scales

**DOI:** 10.1101/2020.09.23.299727

**Authors:** Connie A. Rojas, Santiago A. Ramírez-Barahona, Kay E. Holekamp, Kevin. R. Theis

## Abstract

The gut microbiota is critical for host function. Among mammals, host phylogenetic relatedness and diet are strong drivers of gut microbiota structure, but one factor may be more influential than the other. Here, we used 16S rRNA gene sequencing to determine the relative contributions of host phylogeny and host dietary guild in structuring the gut microbiotas of 11 herbivore species from 5 families living sympatrically in southwest Kenya. Herbivore species were classified as grazers, browsers, or mixed-feeders. We found that gut microbiotas were highly species-specific, and that host taxonomy accounted for more variation in the gut microbiota (30%) than did host dietary guild (10%) or sample month (8%). Overall, similarity among gut microbiotas increased with host phylogenetic relatedness (r=0.73), yet this relationship was not consistent among seven closely related Bovid host species (r=0.21 NS). Within bovids, host dietary guild explained more of the variation in the gut microbiota than did host species. Lastly, while we found that the gut microbiotas of herbivores residing in southwest Kenya converge with those of distinct populations of conspecifics from central Kenya, fine-scale differences in the abundances of bacterial amplicon sequence variants (ASVs) between individuals from the two regions were also observed. Overall, our findings suggest that host phylogeny and taxonomy strongly structure the gut microbiota, but these gut microbial communities could be furthered modified by host ecology, especially among closely related host species.

## Background

The gut microbiota performs critical roles for its host and is essential to host functioning. In mammals, resident gut microbes promote the digestive efficiency of their hosts by synthesizing vitamins, breaking down fiber, and supplementing the host with energy released from fermentation [1–4]. The gut microbiota also interacts with the host immune system, and may also modulate behavior [5–7]. Due to the critical importance of the gut microbiota for host performance, research has focused on determining the forces that shape its assembly and composition. Decades of research show that across vertebrate hosts, the gut microbiota is predominantly shaped by host phylogeny and ecology. Closely related host species tend to have more similar gut microbiotas than more distantly related host species [8–12] and this congruence between host phylogenetic relatedness and gut microbiota similarity is termed “phylosymbiosis” [13–15]. However, gut microbiotas can also be further shaped by their host’s ecology, including their host’s diet, habitat, and geographic location [16–19]. Thus, although both of these host factors may shape the gut microbiota, their relative contributions might be influenced by a variety of variables including the taxonomic breadth of the host species surveyed, and the diversity of host habitats, diets, and physiologies represented.

Similarity in the gut microbiota among individuals could be due to hosts providing potential colonizing microbes with similar ecological niches [20, 21]. This could arise from overlap in diet, habitat, physiology, and behavior among hosts, as well as from shared evolutionary history. Thus, if phylosymbiosis is observed, both host phylogenetic relatedness and ecology could be contributing to the pattern. A diversity of studies have disentangled the effects of these two factors and have shown that phylosymbiosis can be observed among hosts that share habitats or diets, and among hosts that reside in different habitats and consume different diets. For example, in mice, voles, and shrews, gut microbiotas tend to be more similar among closely related host species, despite these animals occupying different habitats [22]. In populations of American pikas (*Ochotona princeps*) from different mountain ranges, a cladogram of gut microbiota similarity was congruent with a phylogeny of host genetic similarity [23]. Among hosts with overlapping diets, gut communities still exhibit patterns consistent with phylosymbiosis, as has been documented for folivorous primates [9]. Furthermore, in 33 species of sympatric herbivores from the Laikipia region in central Kenya, host phylogenetic relatedness strongly predicted gut microbiota composition (r = 0.91) and was weakly correlated with host diet (r = 0.28) [24], suggesting that convergence of gut microbiotas among closely related hosts was not due to similarities in their diet.

Here, we build upon this work and use 16S rRNA gene sequencing to determine the relative influences of host phylogenetic relatedness and host ecology in structuring the gut microbiota of 11 species of herbivores living sympatrically in the Masai Mara National Reserve (henceforth the Masai Mara) in southwestern Kenya. We survey the gut microbiotas of African buffalo, domestic cattle, common eland, impala, Kirk’s dik-dik, Thompson’s gazelle, topi, Masai giraffe, common warthog, plains zebra, and African elephant (Table 1). These species represent 5 mammalian families (Bovidae, Elephantidae, Equidae, Giraffidae, and Suidae) and three dietary guilds: grazers, browsers, and mixed-feeders. Furthermore, we compare the gut microbiotas of conspecific herbivores from the Masai Mara and Laikipia to determine the extent to which host geography and/or local habitat influence gut microbiota composition and patterns of phylosymbiosis. The two regions differ in their altitude, vegetation, mammal densities, and degree of human disturbance [25–29], any of which could potentially affect the gut microbiota compositions of their resident herbivores.

**Table 1.** List of host study species and their associated metadata.

| Order | Family | Species | Dietary Guild | Total Samples (N) | Analyzed samples (N) |
| --- | --- | --- | --- | --- | --- |
| *Cetartiodactyla* | *Bovidae* | African buffalo | grazer | 18 | 17 |
| *Cetartiodactyla* | *Bovidae* | Domestic cattle | grazer | 14 | 13 |
| *Cetartiodactyla* | *Bovidae* | Common eland | mixed feeder | 8 | 8 |
| *Cetartiodactyla* | *Bovidae* | Impala | mixed feeder | 20 | 20 |
| *Cetartiodactyla* | *Bovidae* | Kirk's dik dik | browser | 37 | 31 |
| *Cetartiodactyla* | *Bovidae* | Thomson's gazelle | mixed feeder | 14 | 14 |
| *Cetartiodactyla* | *Bovidae* | Topi | grazer | 19 | 19 |
| *Cetartiodactyla* | *Giraffidae* | Masai giraffe | browser | 25 | 18 |
| *Cetartiodactyla* | *Suidae* | Warthog | mixed feeder | 9 | 8 |
| *Perissodactyla* | *Equidae* | Plains zebra | grazer | 5 | 5 |
| *Proboscidea* | *Elephantidae* | African elephant | mixed feeder | 12 | 12 |

Specifically, we surveyed the gut microbiotas of 11 species of herbivores to 1) determine the relative contributions and amount of variance in the gut microbiota that is explained by host taxonomy (family and species) and host diet, 2) ascertain whether phylosymbiosis is observed across broad host taxonomic scales (i.e. among all study species) and lesser host taxonomic scales (i.e. among 7 closely related Bovid species), and 3) examine the influences of host phylogenetic relatedness and host ecology (diet and geography) on the gut microbiota of conspecific Masai Mara (this study) and Laikipia herbivore hosts [24]. Collectively, our findings elucidate the factors shaping the gut microbiota of hosts at greater and lesser taxonomic scales.

## Results

### Aim 1: Determine the amount of variance in the gut microbiota explained by host taxonomy and host diet

#### Microbiota composition

Our analyses showed that some bacterial taxa were widely shared among host species and dietary guilds, whereas others were abundant only in particular host species. All herbivore gut microbiotas were dominated by two bacterial phyla, *Firmicutes* (51% average relative abundance across samples), and *Bacteroidetes* (32%) (Figure S1). The most abundant bacterial families were *Ruminococcaceae* (30.8%), *Rikenellaceae* (11.4%), *Lachnospiraceae* (10.9%), and *Prevotellaceae* (8%) (Figure 1A). Prevalent bacterial genera included *Alistipes, Bacteroides, Ruminococcus*, and *Treponema* (Figure S2). A total of 10 out of 11, 930 (0.08%) Amplicon Sequence Variants (ASVs) were present in 90% of samples pooled across all host species; 7 were assigned to the family *Ruminococcaceae*, 1 to *Peptococcaceae*, and 2 to *Lachnospiraceae* (*Agathobacter*). According to a blast search against the NCBI nucleotide database, sequences from the 7 *Ruminococcaceae* ASVs were highly similar to sequences from uncultured *Ruminococcaceae* strains, uncultured rumen bacteria, and uncultured anaerobic bacteria. Of these 10 ASVs, only two were abundant across samples (e.g. ASV10543 *Ruminococcaceae*), five were modestly abundant in specific host species (e.g. ASV71 *Agathobacter* in elephants), and 3 were present at very low abundances in all samples (e.g. ASV7824 *Peptococcaceae*) (Figure S3). The latter 3 ASVs do not appear to be contamination introduced during DNA extraction and sequencing, as these sequences are highly similar to those found in rumen and fecal samples.

**Figure 1.**
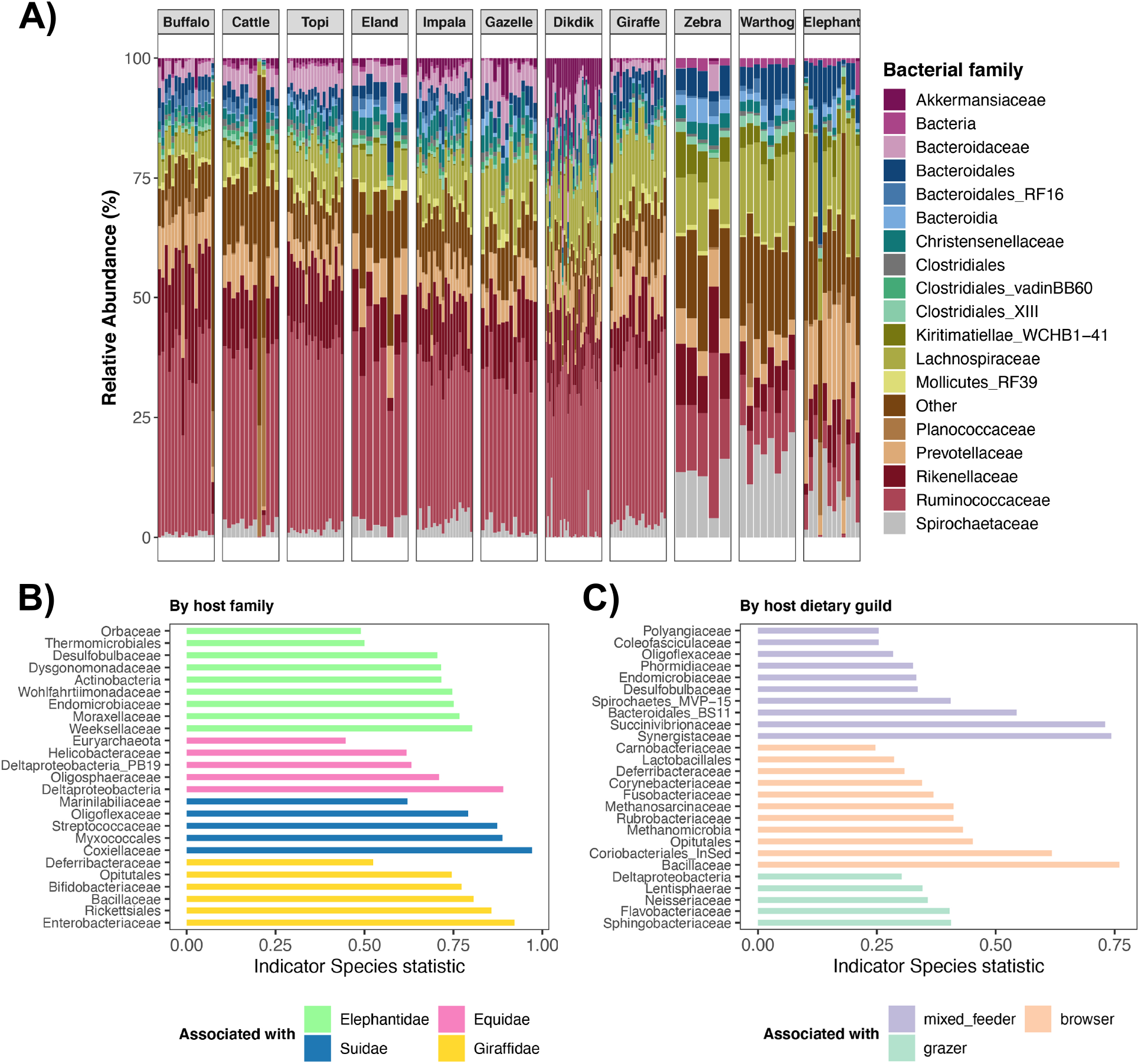
Gut microbiota composition of African herbivores. **A)** Stacked bar plots showing the relative frequency of 16S rRNA gene sequences assigned to each bacterial family (or order, if a family-level classification could not be assigned) across samples. Samples are grouped by host species, and each color represents a bacterial family. **B)** Bacterial families significantly associated with particular herbivore families as determined by indicator species analysis. Differences in these taxa abundances can explain differences in the microbiota among the different groups. **C)** Bacterial families significantly associated with herbivores from different dietary guilds as determined by indicator species analysis.

Nonetheless, variation in gut microbiota compositions among host species and dietary guilds was clearly evident. Indicator species analysis showed that the gut microbiotas of elephants were significantly associated with *Endomicrobiaceae* and *Desulfobulbaceae*, those of Zebras with *Helicobacteraceae* and *Deltaproteobacteria*, and those of warthogs with *Myxococcales* and *Coxiellaceae* (Figure 1B). Giraffe gut microbiotas were highly associated with *Enterobacteriaceae, Bifidobacteriaceae*, and *Bacillaceae*. Similarly, there were bacterial types that were indicative of specific dietary guilds. Grazer gut microbiotas were characterized by *Sphingobacteriaceae, Flavobacteriaceae, Neisseriaceae*, and *Lentisphaeria* (Figure 1C). Browser gut microbiotas had 11 indicator bacterial taxa, including *Bacillaceae, Coriobacteriales, Methanomicrobia* and *Rubrobacteriaceae*. Lastly, the gut microbiotas of mixed feeders were highly associated with *Synergistaceae, Succinivibrionaceae*, and *Bacteroidales*, among other bacteria.

More fine-scale analysis of the presence and absence of bacterial ASVs also revealed that the gut microbiotas of our studied herbivores contained microbes that were biased towards particular host species. These were bacterial ASVs that were present in 75% samples for that host species, and absent in 97% of other samples. Buffalo, cattle, topi, and impala gut microbiotas mostly contained ASVs that were present in other herbivores, as <3% of their ASVs were biased to any of these host species. Between 4% and 8% of ASVs comprising the gut microbiota of dik-diks, eland, elephant, Thompson’s gazelle, and giraffe were biased towards these host species. Warthogs and zebras however, harbored more unusual microbiotas, as 70-77% of their ASVs were rarely detected in the guts of the other African mammals.

#### Microbiota alpha-diversity

Gut microbiota richness, evenness, and phylogenetic diversity also varied with host taxonomy (family and species) and dietary guild (Table 2, Figure 2). Post-hoc comparisons revealed that hosts from the Suidae and Elephantidae families generally harbored less rich and less even gut communities than the other surveyed host families (Figure 2A; Table S1). Moreover, equids harbored more phylogenetically diverse gut communities than all other herbivores (Table S1). Across the three alpha-diversity metrics, browsers had less diverse gut microbiotas than grazers or mixed-feeders (Figure 2A, Table S2). Within a single host family (i.e. Bovidae), the gut microbiotas of buffalo, dikdiks, and gazelles were less rich, even, and phylogenetically diverse than those of all other sampled bovids (Figure 2B, Table S3). In bovid hosts, browsers also had less diverse gut microbiotas than grazers or mixed-feeders (Figure 2B, Table S3).

**Table 2.**
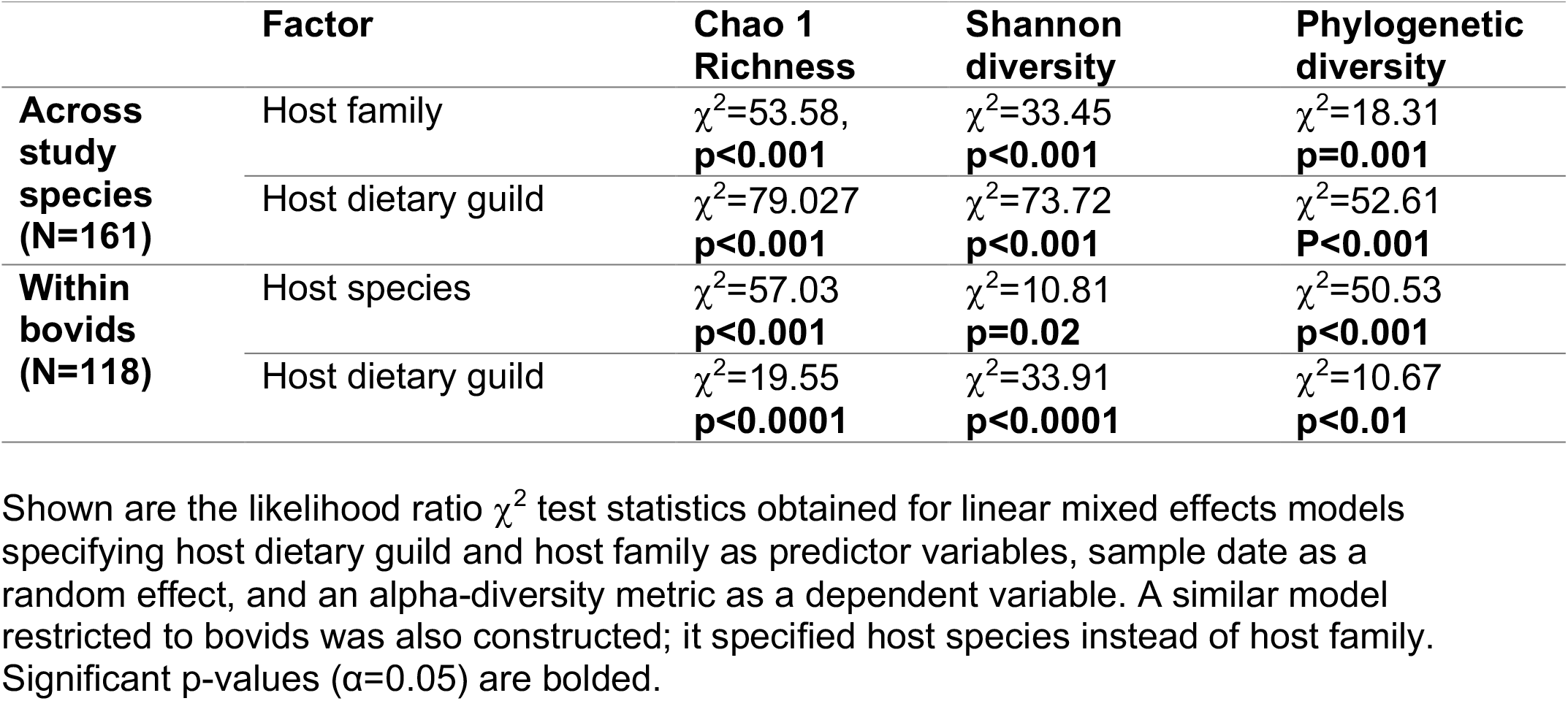
Microbiota richness, evenness, and phylogenetic diversity vary with host taxonomy and dietary guild.

|  | Factor | Chao 1 Richness | Shannon diversity | Phylogenetic diversity |
| --- | --- | --- | --- | --- |
| **Across study species (N=161)** | Host family | $\chi^2=53.58$,<br>**p<0.001** | $\chi^2=33.45$<br>**p<0.001** | $\chi^2=18.31$<br>**p=0.001** |
| | Host dietary guild | $\chi^2=79.027$<br>**p<0.001** | $\chi^2=73.72$<br>**p<0.001** | $\chi^2=52.61$<br>**P<0.001** |
| **Within bovids (N=118)** | Host species | $\chi^2=57.03$<br>**p<0.001** | $\chi^2=10.81$<br>**p=0.02** | $\chi^2=50.53$<br>**p<0.001** |
| | Host dietary guild | $\chi^2=19.55$<br>**p<0.0001** | $\chi^2=33.91$<br>**p<0.0001** | $\chi^2=10.67$<br>**p<0.01** |

Shown are the likelihood ratio $\chi^2$ test statistics obtained for linear mixed effects models specifying host dietary guild and host family as predictor variables, sample date as a random effect, and an alpha-diversity metric as a dependent variable. A similar model restricted to bovids was also constructed; it specified host species instead of host family. Significant p-values ( $\alpha=0.05$ ) are bolded.

**Figure 2.**
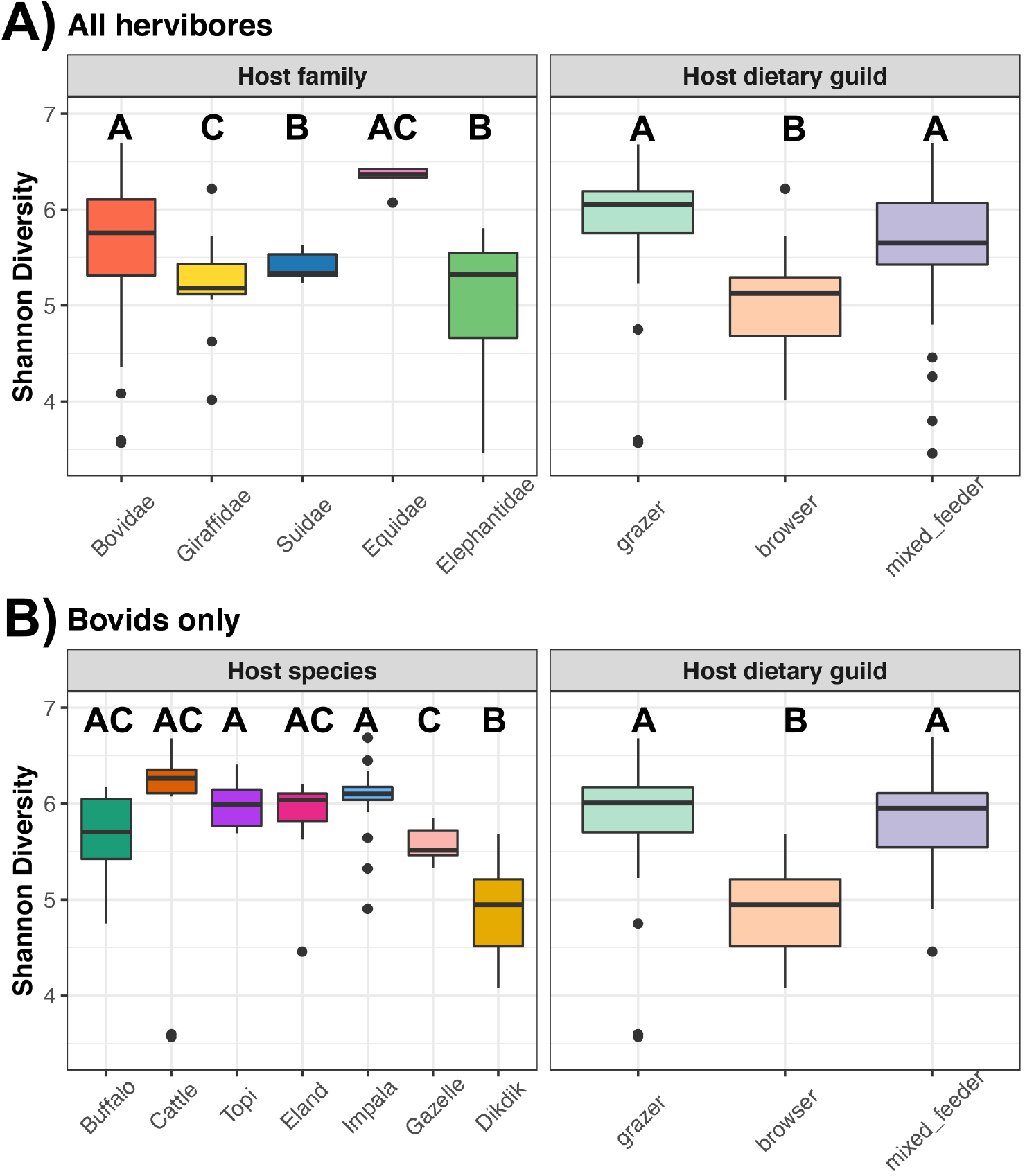
Host taxonomy and dietary guild are associated with gut microbiota diversity in African herbivores. **A)**Boxplots of microbiota evenness (Shannon diversity) among host families and dietary guilds across all studied herbivores, and **B)** among host species and dietary guilds within the family *Bovidae*. Boxes that do not share any letters represent statistically significant comparisons; see Tables S1-S3 for all post-hoc comparisons. Thicker dots represent outlier values.

Additionally, we determined whether *similarity* in gut microbiota alpha-diversity was correlated with a *quantitative measure* of host diet, while accounting for variation in the gut microbiota associated with host phylogenetic relatedness. Specifically, we used previously published dietary C4 (%) data [30–33] from these 11 herbivore species as our numerical measure of diet (Table S4). C4 (%) values quantify the degree of grazing by the host and reflect the proportion of an animal’s diet composed of C4 monocotyledon grasses relative to C3 trees, shrubs, and forbs. Similarity in gut microbiota *evenness* (Shannon diversity) increased with host overlap in dietary C4, but this was not true of gut microbiota richness or phylogenetic diversity (partial Mantel Chao1 r=0.03, p=0.34; Shannon r=0.35, p=0.028, PD r=0.097, p=0.25).

#### Microbiota beta-diversity

Across the surveyed herbivores, host family explained on average ∼23% of the variation in gut microbiota structure, followed by host dietary guild (10%), sample month (8%), and host species (7.5%) (PERMANOVA analyses, Table 3). Regardless of whether distance matrices took into account the proportions of bacterial taxa, their presence/absence, or their phylogenetic relatedness, the percent variation explained by each host factor was consistent. Therefore, for brevity, we only present PCoA ordination plots using the Bray-Curtis index. These plots show that gut microbiotas primarily partition by host family, and also secondarily by host dietary guild (Figure 3A).

**Table 3.** Host taxonomy and dietary guild shape the gut microbiotas of African herbivores.

| **Analysis** | **Host factors** | **Bray-Curtis**<br>**(% variance explained)** | **Jaccard**<br>**(% variance explained)** | **Weighted**<br>**Unifrac**<br>**(% variance explained)** | **Unweighted**<br>**Unifrac**<br>**(% variance explained)** |
| --- | --- | --- | --- | --- | --- |
| **Across all host study species (N=165)** | Host family | 22.33,<br>**p=0.001** | 20.59,<br>**p=0.001** | 26.38,<br>**p=0.001** | 24.23,<br>**p=0.001** |
|  | Host dietary guild | 11.25,<br>**p=0.001** | 10.21,<br>**p=0.001** | 9.71,<br>**p=0.001** | 10.30,<br>**p=0.001** |
|  | Host species | 9.40,<br>**p=0.001** | 8.6,<br>**p=0.001** | 4.45,<br>**p=0.001** | 7.23,<br>**p=0.001** |
|  | sample month | 7.42,<br>**p=0.001** | 6.80,<br>**p=0.001** | 10.21,<br>**p=0.001** | 7.66,<br>**p=0.001** |
| **Across bovids (N=122)** | Host dietary guild | 18.26,<br>**p=0.001** | 15.91,<br>**p=0.001** | 18.35,<br>**p=0.001** | 16.77,<br>**p=0.001** |
|  | Host species | 15.16,<br>**p=0.001** | 13.38,<br>**p=0.001** | 8.23,<br>**p=0.001** | 12.16,<br>**p=0.001** |
|  | sample month | 7.56,<br>**p=0.001** | 7.03,<br>**p=0.001** | 7.59,<br>**p=0.001** | 7.17,<br>**p=0.001** |

Shown are the $R^2$ values (% variance explained) and p-values for PERMANOVA tests ( $y \sim$ sample month + host dietary guild + host taxonomy) based on 4 types of distance matrices. Bray-Curtis and Weighted Unifrac distance matrices take into consideration the proportions of bacterial taxa, while Jaccard and unweighted Unifrac take into account only their presence or absence. Both Unifrac distances account for phylogenetic relatedness among bacterial types. Significant p-values ( $\alpha=0.05$ ) are bolded.

**Figure 3.**
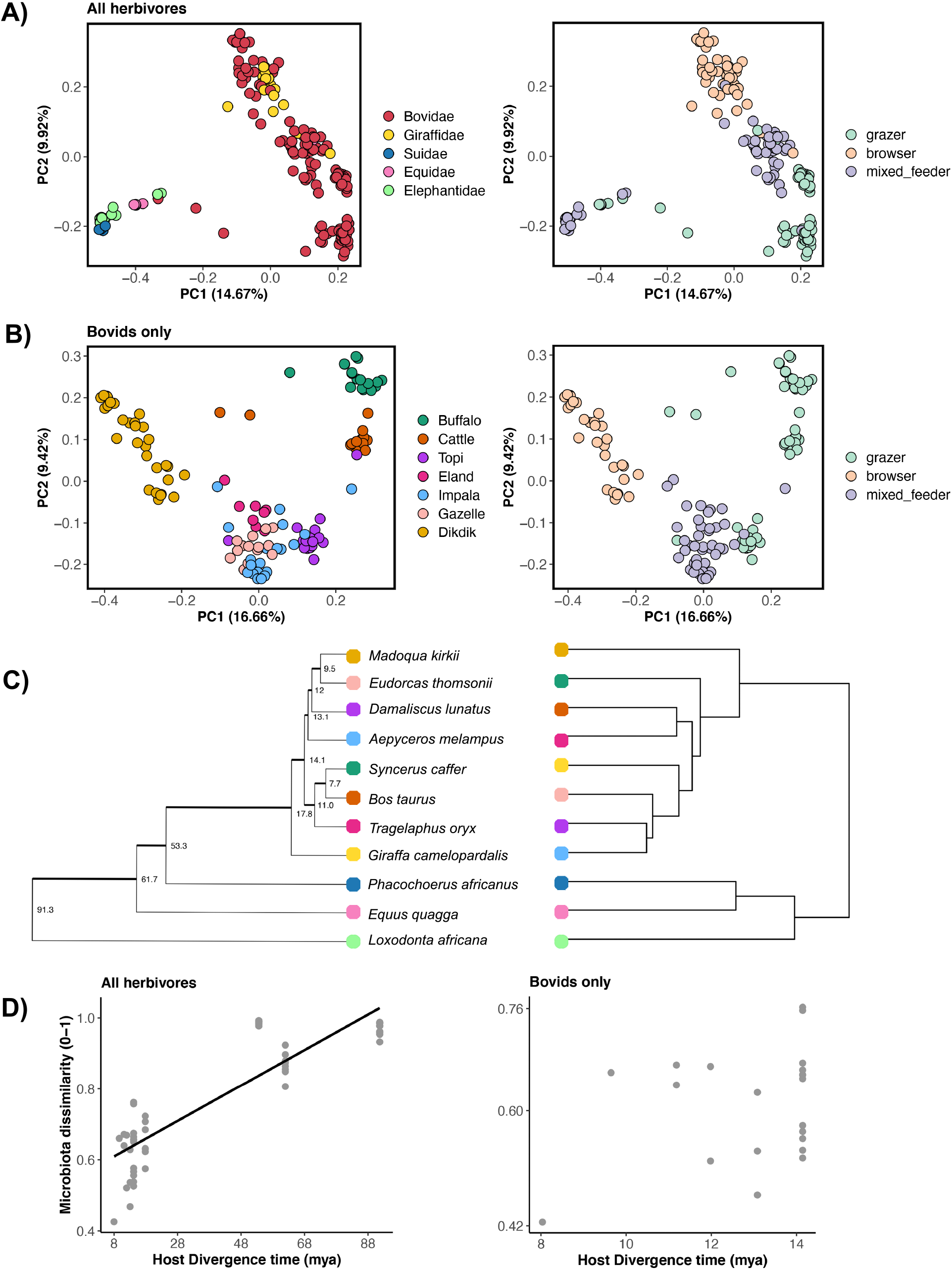
African herbivore gut microbiotas exhibit patterns consistent with phylosymbiosis. **A)** PCoA plots constructed from Bray-Curtis dissimilarity matrices. Each point represents a sample and is color-coded by host family (left) or host dietary guild (right). Closeness of points indicates high community similarity. The percentage of variance accounted for by each principal-coordinate axis is shown in the axis labels. **B)** PCoA plots constructed from Bray-Curtis dissimilarity matrices of bovid species only. Each point is color-coded by host species (left) or host dietary guild (right). **C)** Phylogenetic tree of host species (left) obtained from pruning Upham’s *et al*. 2019 Mammalian supertree, compared against a dendogram (right) of gut microbiota similarity using hierarchical clustering. **D)** Scatterplot of pairwise host divergence times (in millions of years) vs. gut microbiota similarity (Bray-Curtis distances) across all sampled herbivores (left) and within the single host family *Bovidae* (right).

Within Bovidae, host dietary guild was a slightly stronger predictor of the gut microbiota than host species or sample month. On average, host dietary guild accounted for 17.3% of the variation, whereas host species explained 12.2% of the variation, and sample month contributed to 7.3% of the variation (PERMANOVA analyses, Table 3). PCoA ordinations showed that the gut microbiotas of bovids clustered by host dietary guild and host species (Figure 3B). When controlling for host dietary guild and sample month, the microbiota is highly-species specific and host species accounts for 40% of the observed variation across samples (PERMANOVA: Bray-Curtis R^2^=0.42; Jaccard R^2^=0.39; Weighted Unifrac R^2^=0.40, Unweighted Unifrac R^2^=0.41; all p=0.001) (Figure S4).

### Aim 2: Ascertain whether phylosymbiosis is observed among herbivore hosts at broad and lesser taxonomic scales

Although a dendrogram of gut microbiota similarity did not closely reflect host phylogeny (Figure 3C), gut microbiota similarity did increase overall with host phylogenetic relatedness among the 11 species of herbivores (mantel test Bray-Curtis r=0.76, p=0.006; Jaccard r=0.72, p=0.006; Weighted Unifrac r=0.75, p=0.007; Unweighted Unifrac r=0.72, p=0.007) (Figure 3D). The gut microbiotas tended to be more similar among closely related host taxa (e.g. buffalo and cattle) than among distantly related host taxa (e.g. impala and elephant). Despite warthogs being more closely related to bovids than to elephants and zebras, they had highly dissimilar communities from all of the herbivore species examined, and pairwise comparisons that include this host species deviate from the trendline (∼1.0 Bray-Curtis dissimilarity, Figure 3D).

Importantly, at a lesser host taxonomic scale, among closely related host species in the Bovid family, we did not find a consistent relationship between gut microbiota similarity and host phylogenetic relatedness (mantel test Bray-Curtis r=0.27, p=0.09; Jaccard r=0.20, p=0.08; Weighted Unifrac r=0.22, p=0.059; Unweighted Unifrac r=0.16, p=0.08) (Figure 3D). However, when taking into account fine-scale variation associated with host diet (%C4) (Table S4), gut microbiota similarity is modestly correlated with host phylogenetic relatedness (partial Mantel Bray-Curtis r=0.43, p=0.013; Jaccard r=0.35, p=0.016; Unifrac r=0.33, p=0.017; Unweighted Unifrac r=0.29, p=0.017). When repeating this analysis on the dataset that includes all herbivore hosts, the original finding remained unchanged; accounting for fine-scale variation in host %C4 does not affect the phylosymbiosis pattern observed at a broad host taxonomic scale (partial Mantel Bray-Curtis r=0.77, p=0.002, Jaccard r=0.72, p=0.007, Unifrac r=0.75, p=0.004, Unweighted Unifrac r=0.72, p=0.01). In summary, we observed phylosymbiosis among hosts spanning a broad taxonomic scale involving multiple families, but not consistently among hosts spanning a lesser taxonomic scale involving only members of the Bovid family.

### Aim 3: Examine influences of host phylogenetic relatedness and host ecology on the gut microbiota of conspecific hosts

Lastly, we further analyzed the influences of host phylogenetic relatedness and host ecology (diet and geography) on the gut microbiota of 8 herbivore species from two distinct populations in the Masai Mara (this study) and the Laikipia region, respectively (Kartzinel *et al*. [24]). The eight herbivore species overlapping both studies were African buffalo, domestic cattle, common eland, impala, giraffe, plains zebra, common warthog, and African elephant.

We found that gut microbiota structure varied little among hosts from the two geographic regions, as this factor accounted for <3% of the variation in the gut microbiota (PERMANOVA analyses, Table S5). The gut microbiotas were primarily structured by host species and host dietary guild, which explained on average, 38% and 11% of the variation, respectively (PERMANOVA analyses, Table S5); sample month explained an additional 7% of the variation. Ordination plots confirm these findings and demonstrate that samples primarily cluster by host species (Figure 4A), although some separation of samples based on host geographic region is also observed, particularly among cattle, impala, and giraffe.

**Figure 4.**
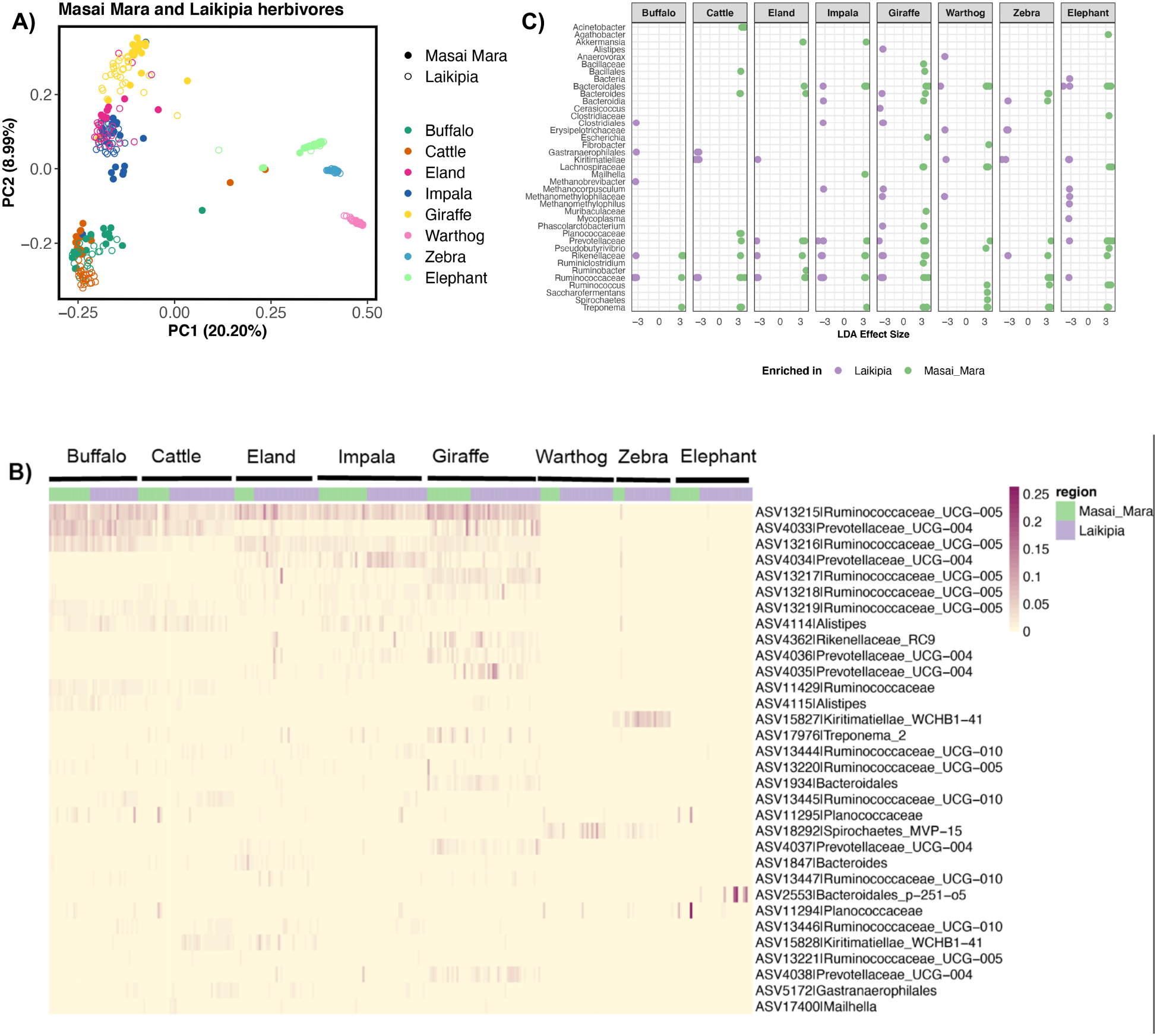
The gut microbiotas of conspecific African herbivores broadly converge, but also exhibit differences in their ASV abundances. We compared the gut microbiotas of eight species of herbivores residing in both the Masai Mara (this study) and Laikipia (Kartzinel *et al*. 2019) regions in Kenya. **A)** PCoA plot constructed from Bray-Curtis dissimilarity matrices. Each point represents a sample and is color-coded by host species; shape shading indicates geographic region. **B)** Heatmap of the 32 most abundant bacterial ASVs residing in the gut microbiotas of Masai Mara and Laikipia herbivores. Samples are grouped by host species, and are color-coded by host geographic region. **C)** ASVs enriched in Masai Mara vs. Laikipia herbivores as determined by LEfSe. Each dot represents a unique ASV and is color-coded by host geographic region. A total of 210 ASVs are displayed (those with LDA >3.2) and their family or genus level classification are on the x- axis.

For this dataset, a total of 3 ASVs out of 18, 039 (0.01%) were present across 90% of samples; two were classified as *Ruminococcaceae*, and 1 as *Lachnospiraceae*. All 3 ASVs were among the 10 ASVs present across 90% of Masai Mara samples. To further compare the gut microbiotas of conspecific hosts, we analyzed the relative abundances of the 32 most abundant ASVs in the dataset. A heatmap of these 32 ASVs demonstrate that there were ASVs that were found at comparable proportions in conspecific hosts, as well as ASVs that were differentially abundant among conspecific hosts (Figure 4B, Table S6). For example, ASV4033 *Prevotellaceae* is similarly abundant between Masai Mara and Laikipia hosts, but ASV15828 *Kiritimatiellae* appears to be enriched in cattle from Laikipia compared to cattle from the Masai Mara. Linear mixed models that statistically compared the relative abundances of these ASVs between the two groups showed that 13 of the 32 ASVs were actually differentially abundant between conspecifics (Table S6).

To extend these analyses and identify additional ASVs that may be differentially abundant among conspecifics, we also conducted Linear discriminant analysis Effect Size (LEfSe) for each host species. Roughly 30% of the ASVs in the gut microbiotas of each host species were enriched in hosts from one population relative to the other (Table S7). ASVs that were typically enriched were classified as *Ruminococcaceae, Lachnospiraceae, Rikenellaceae, Clostridiales, Bacteroidales*, and *Kiritimatiellae* (Figure 4C). Herbivores from Laikipia tended to be enriched in *Methanocorpusculum, Clostridiales*, and *Kiritimatiellae* ASVs relative to Masai Mara herbivores (Figure 4C). Masai Mara gut microbiotas were overrepresented by *Lachnospiraceae* and *Treponema* ASVs. Interestingly, hosts from both geographic regions could be enriched in taxonomically similar ASVs. For example, eland in Laikipia were enriched in 3 ASVs classified as *Prevotellaceae, Rikenellacea*e, and *Ruminococcaceae*, respectively, and eland from the Masai Mara were enriched in 3 different ASVs that were also classified as *Prevotellaceae, Rikenellacea*e, and *Ruminococcaceae* (Figure 4C). These findings suggest that variation in the gut microbiotas of these herbivore conspecifics is observable at the level of specificity of bacterial ASVs.

Patterns consistent with phylosymbiosis were also observed in this combined dataset, despite herbivore hosts occupying different geographic regions in Kenya, and representing distinct populations. Gut microbiota similarity increased with host phylogenetic relatedness (mantel test Bray-Curtis r=0.60, p=0.03; Jaccard r=0.64, p=0.04; Weighted Unifrac r=0.54, p=0.003; Unweighted Unifrac r=0.66, p=0.04), although the strength of the correlation was less than that of Masai Mara herbivores only (r=0.76), or what was previously reported for Laikipia herbivores (r=0.91) [24]. Consistent with our earlier findings, phylosymbiosis was not evident among the four species of bovids that overlapped between the two studies (mantel test Bray-Curtis r=0.30, p=0.25; Jaccard r=0, p=0.5; Weighted Unifrac r=-0.52, p=0.91; Unweighted Unifrac r=-0.49, p=0.83). When taking into account gut microbiota variation attributable to host diet (% dietary C4), both findings remained unchanged (<u>partial Mantel test all herbivores</u>: Bray-Curtis r=0.60, p=0.03, Jaccard r=0.64, p=0.04, Weighted Unifrac r=0.53, p=0.03, Unweighted Unifrac r=0.65, p=0.05 0.05; <u>bovids only</u>: Bray-Curtis r=0.36, p=0.16, Jaccard r=0, p=0.5, Weighted Unifrac r=-0.52, p=0.87, Unweighted Unifrac r=-0.61, p=0.83).

## Discussion

### Principal findings of study

The primary purpose of this study was to determine the relative contributions of host phylogenetic relatedness and dietary guild in structuring the gut microbiotas of 11 species of sympatric African herbivores. We also compared the gut microbiotas of herbivores from the Masai Mara to herbivores from Laikipia, Kenya to determine the extent to which two distinct populations of identical herbivore species varied in their gut microbiotas. We found that gut microbiotas were highly species-specific, but also varied with host ecology, including host dietary guild and sample month, particularly in closely related Bovid species. Furthermore, gut microbiota similarity increased with host phylogenetic relatedness at broad host taxonomic scales, but at lesser taxonomic scales, the phylosymbiosis signal was only apparent after accounting for fine-scale variation attributable to host diet (%C4). Lastly, although the gut microbiotas of conspecific herbivore hosts converged and primarily clustered by host species, differences between conspecifics in the relative abundances of their bacterial ASVs were also observed. Collectively, our findings suggest that mammalian gut microbiotas are strongly shaped by host phylogenetic relatedness and taxonomy, but they can be further modified by host ecology, including host diet and geography.

### Aim 1: Determine the amount of variance in the gut microbiota explained by host taxonomy and host diet

Across the surveyed herbivores, gut microbiota composition, diversity, and structured varied with host taxonomy. Species-specificity of the gut microbiota is widespread, and is commonly reported in the majority of comparative gut microbiome studies [11, 34–36]. Host species may vary in their body size, behavior, neuroendocrine system, immune function, and metabolism, any of which could potentially shape the structure of their gut microbiotas [37–40]. When comparing gut microbiota alpha-diversity, results showed that warthogs and elephants harbored less diverse gut communities than the other sampled herbivores. Due to their omnivory, warthogs have a greater dietary breadth than the other studied herbivores, yet they harbored less diverse microbiotas. This is in accordance with prior findings, which report that the most diverse diets do not always correlate with the most diverse gut microbiotas [24, 41, 42]. Furthermore, analyses showed that browsers had the least diverse gut microbiotas, potentially because they consume vegetation that has a higher lignin content and a lower fiber digestibility than grass [43]. Specialized bacterial metabolisms may be required to digest this tougher plant material. Additionally, group size has been shown to correlate with gut microbiota diversity [44, 45], and the browsers in our study (giraffes, dikdiks) forage in smaller groups than grazers (buffalo, zebras) and mixed-feeders (gazelles, impala), which forage in herds. Frequent social interactions and interactions with a greater number of individuals is known to promote species richness in individual gut microbiotas [45, 46].

Similar to findings from a plethora of microbiome studies, the gut microbiota structure of the studied herbivores also varied with host diet. Hosts from each dietary guild consume food sources that vary in their structure, chemistry, and nutrition quality; which require morphological, physiological and behavioral adaptations [47, 48]. For example, grazers mostly feed on grasses, which have thicker cell walls, a lower protein content, and use the C4 photosynthetic pathway compared to browse (e.g. leaves, shrubs, and woody vegetation), which have a higher protein content, and use C3 photosynthesis [47, 48]. To efficiently extract energy from these different food sources, browsers and grazers evolved adaptations in their salivary chemistry, tooth morphology, gut structure, and speed of digestion [49, 50]. These adaptations, along with the actual nutrients hosts are providing to their microbes, potentially contribute to gut microbiota divergence among hosts from different dietary guilds.

Despite differences in the gut microbiota among host species and dietary guilds, there were some features of the gut microbiota that were shared across individuals from multiple species. Across our surveyed herbivores, the most abundant bacterial taxa in the gut microbiota were *Ruminococcaceae, Rikenellaceae, Lachnospiraceae*, and *Prevotellaceae* which represent core taxa previously found in the gut microbiotas of many ruminants and herbivores in general, including cervids and bovids [8, 51], equids [52], elephants [53], and giraffes [54]. *Ruminococcaceae* and *Lachnospiraceae* have also been found in the guts of folivorous primates [3] and in domestic pigs [55, 56]. Members of these bacterial families are responsible for digesting the cellulose, hemicellulose, lignin, and protein found in vegetation, and fermenting these into short-chain fatty acids (SCFAs) [57]. SCFAs represent usable forms of energy for the hosts [58] and contribute to host colonocyte growth, immune defense, and anti-inflammatory responses [1]. These bacterial taxa also possess fiber-degrading capabilities and can provide their hosts with protection against ingested toxic plant secondary metabolites [59].

Interestingly, 7 of the 10 ASVs that were present in 90% of Masai Mara herbivores were classified as *Ruminococcaceae* and were sequences highly similar to uncultured *Ruminococcaceae* strains extracted from bovine, goat, and sheep rumens [60], suggesting that these “core” microbes could be functionally important for the host.

### Aim 2: Ascertain whether phylosymbiosis is observed among herbivore hosts at broad and lesser taxonomic scales

Our results showed that phylosymbiosis was observed across a broad host taxonomic scale, among 11 species of herbivores living sympatrically in the Masai Mara. Patterns of phylosymbiosis have been documented extensively in many vertebrate groups, including primates, rodents, ruminants, carnivores, reptiles, and insects [9–12, 24, 61, 62]. Evidence of phylosymbiosis among host species living in *sympatry* specifically, has been previously documented in seven species of deer mice [63], six species of Malagasy mammals [64], twelve species of lemurs [65], and nine species of diurnal, non-human primates [66].

The mechanisms and processes that yield patterns of phylosymbiosis have not yet been elucidated, but host ecological and phenotypic traits are likely acting as filters and thus shaping microbial community assembly. Closely related hosts are potentially colonized by taxonomically similar microbes due to similarities in their morphology, anatomy, digestive physiologies, and immune system components [37, 38, 67]. Specifically, related hosts may possess similar antimicrobial peptides and toll-like receptors that serve to filter the same bacterial clades from the environment [68, 69]. Closely related hosts may further develop immune tolerance via adaptive immunity to the same symbiotic, commensal, and transient microbes [68, 69]. Lastly, some phylogenetically related hosts may also possess similar social group structures and pathways for transmitting microbes among group-mates, thereby contributing to patterns of phylosymbiosis. Overall, accumulation of differences in traits as hosts diverged from one another could potentially provide enough niche differentiation in the gut to promote the divergence of their symbiotic bacterial communities.

At a lesser host taxonomic scale, within our sampled group of closely related Bovid species in the Masai Mara, variation in the gut microbiota was more strongly associated with host diet than host species, and a pattern consistent with phylosymbiosis was only apparent if fine-scale variation in diet (%C4) was accounted for. This phylosymbiosis signal was weaker (r=0.35) than that observed for all herbivores (r=0.73). Similarly, other studies report that among closely related hosts, host ecology more strongly predicts the structure of the gut microbiota than host relatedness. For example, in lemurs (*Eulemur* spp., *Propithecus* spp.), phylosymbiosis was observed across but not within two host lineages, and within host lineages, host habitat (dry forest vs. rainforest) was significantly correlated with gut microbiota diversity [65]. In populations of yellow (*Papio cynecephalus*) and anubis baboons (*Papio anubis*), gut microbiota dissimilarity did not increase with host genetic distance, but did vary with their habitat’s soil chemistry [70]. Because the bovids surveyed here are closely related, their gut microbiotas are already very similar, and variation can result from fine-scale differences in diet (proportions of grass vs. shrubs vs. trees consumed) [43, 71–73]. Nonetheless, some of the variation in the gut microbiota of bovids is not attributable to host diet and can be explained by host phylogenetic relatedness. Thus, even among closely related host species, both ecological and evolutionary forces shape gut microbial communities.

### Aim 3: Examine influences of host phylogenetic relatedness and host ecology on the gut microbiota of conspecific hosts

While gut microbiota structure was primarily associated with host species in the combined Masai Mara and Laikipia dataset, differences in gut microbiota composition between conspecific hosts from the two populations were also evident. Herbivores of the same species may possess similar evolutionary trajectories, physiologies, and behaviors, and thus may be providing microbes with similar niches for colonization, which is why at a broad level their gut microbiotas converge. However, the two geographic regions do vary in their climate, soil geochemistry, plant communities, and resident herbivore species [25–29, 74], and potentially in their bacterial species pools, which could lead to the fine-scale microbiota differences among conspecifics. This finding was supported by our data; over 30% of ASVs were differentially enriched between Masai Mara and Laikipia hosts. Laikipia herbivores for example, tended to be enriched in ASVs classified as *Methanocorpusculum, Clostridiales*, and *Kiritimatiellae*, while Masai Mara herbivores had an overrepresentation of ASVs belonging to *Lachnospiraceae* and *Treponema*. Abundances of the methanogenic *Methanocorpusculum* are related to forage type and geographic location in cattle [75], and in the mammalian gut, *Lachnospiraceae* are associated with a high-fat diet [76]. Furthermore, in the bovine rumen, *Treponema* degrade hemicellulose and their growth increases in the presence of pectin [77, 78], a carbohydrate abundant in non-woody plants. This suggests that differences in the ASV abundances between conspecific hosts likely reflect fine-scale differences in their diets and habitats. Interestingly, both *Clostridiales* and *Lachnospiraceae* are major microbial taxa of the mammalian gut and comprise fermentative bacteria that synthesize SCFAs from the hydrolysis of starches and sugars [57]; thus, conspecific hosts can be enriched in taxonomically distinct microbes that perform similar functions.

Notably, 3 “core” bacterial ASVs were present in > 90% of samples from the 8 species of herbivores sampled from two allopatric populations. The 3 ASVs were highly similar to sequences from uncultured *Ruminococcaceae* strains and other anaerobic, rumen bacteria, illustrating that these ASVs a) may be easily acquired from the environment, and b) could be essential for host digestion.

Lastly, we found that phylosymbiosis was also evident among conspecific African herbivores living in allopatry, although the strength of the phylosymbiotic signal was slightly reduced compared to that observed for either sympatric population considered in isolation. Overlap in gut microbiota structure is thought to be lower in allopatric animal populations than in sympatric animals due to variation introduced by habitat, dietary differences, and the spatial limits of bacterial dispersal [12]. It is important to note that differences between the gut microbiotas of Laikipia and Masai Mara conspecifics could also be potentially attributable to differences in sampling, DNA extraction, and sequencing protocols between the two studies [79–81]. Collectively, our findings show that mammalian gut microbiotas converge among closely related host species and among conspecifics, but can be differentiated with variation introduced by the host’s ecology.

## Conclusions

Our study showed that among 11 species of African herbivores living in sympatry, gut microbiotas are highly species-specific and exhibit patterns congruent with phylosymbiosis. However, these gut microbiotas are also shaped by their host’s ecology, and we observed differences between hosts from different dietary guilds Furthermore, among eight species of herbivores residing in two geographic regions in Kenya, gut microbiotas were similar among hosts of the same species, but also exhibited fine scale differences in the abundances of their bacterial ASVs. Overall, these findings suggest that related hosts are providing microbes with similar niches for colonization, but these microbial niches are further shaped by host diet, geography, and local environmental conditions. Future studies should examine whether phylosymbiosis is evident at the functional level in the gut metagenomes of African herbivores and in metazoan taxa in general. The gut microbiotas between distantly related host species could be taxonomically dissimilar, but functionally similar due to the widespread functional redundancy observed in microbial metabolisms. These studies will be necessary to further our understanding of the processes and mechanisms potentially underlying patterns of phylosymbiosis.

## Methods

### Study location and sampling

Fecal samples (N=181) were collected opportunistically from 11 species of herbivores permanently residing in the Talek and Mara Triangle regions of the Masai Mara (1°22′19″S, 34° 56′17″E) from March-June 2018 (Table 1). This Reserve is covered by open rolling grassland interspersed with seasonal watercourses and riparian vegetation. It has two rainy seasons (March-May and November-December, with annual rainfall >1000mm) [82], and most of our sampling took place during the rainy months, particularly during the month of March (Figure S5). Although the Masai Mara is home to small resident populations of zebra and wildebeest, millions of these individuals migrate into the Reserve from July-October every year. Because our sampling occurred before July, samples from wildebeest and zebras were limited.

For fecal sample collection, we either observed animals defecating or identified species-of-origin based on the size, shape, and consistency of fresh dung, following Kartzinel *et al*. [24]. Samples were then placed in sterile cryogenic vials and stored in liquid nitrogen until they were transported to Michigan State University, where they remained frozen at -80°C until nucleic acid extraction. For a list of samples and their associated metadata, see the Github repository for this project (https://github.com/rojascon/Rojas_et_al_2020_African_herbivores_gut_microbiome).

While we did not directly collect diet data from the surveyed herbivores, we used Kingdon’s *East African Mammals* [83–86] to classify our study species into grazers, browsers, and mixed-feeders. To obtain more fine-scale data on host diet, we also compiled dietary C4 (%) data for these herbivores from previously published studies (Table S4) [30– 33]. Percent C4 values reflect the proportion of monocotyledon grasses consumed relative to trees, shrubs, and forbs.

### DNA extraction and 16S rRNA gene sequencing

Fecal samples were sent to the University of Chicago at Illinois (UIC) Sequencing Core for automated DNA extractions using QIAGEN DNeasy PowerSoil kits (Valencia, CA, USA). DNA concentrations of the fecal sample extracts were quantified using Qubit. The V4 region of the 16S rRNA gene was targeted for sequencing on the Illumina MiSeq platform at the Michigan State University Genomics Core, using published protocols by Caporaso *et al*. 2012 [87] and Kozich *et al*. 2013 [88].

### Sequence processing and bioinformatics

Sequences from Masai Mara herbivore gut microbiotas were processed in R (v.3.6.2) [89] using the Divisive Amplicon Denoising Algorithm (DADA2) pipeline (v.1.14.1) [90] to infer amplicon sequence variants (ASVs). Briefly, reads were filtered for quality, allowing for 2 and 3 errors per forward and reverse read, respectively (trimLeft = c(10, 10), maxN = 0, maxEE = 2, truncQ = 2). Forward reads were trimmed to 240bp and reverse reads to 200bp; these paired-end reads were merged. Sequences were then dereplicated to remove redundancy and ASVs were inferred by pooling reads from all samples. Prior to creating the ASV abundance table, chimeras were removed and ASVs were taxonomically classified using the SILVA rRNA gene reference database (v.132) [91] with an 80% confidence threshold. ASVs taxonomically assigned as Eukarya, Chloroplasts, or Mitochondria were removed from the dataset, as were those of unknown Kingdom origin; 12,938 total ASVs remained. The resulting ASV table and the taxonomic designations of the ASVs are available on GitHub. On average, samples retained over 70% (± 11%) of their total sequences after processing in DADA2. Nineteen samples did not amplify well (<400 sequences after processing) and were removed from the dataset. Most of these samples belonged to browser species (giraffes and dik-diks) suggesting that there may have been PCR inhibitors in their fecal samples (e.g. humic acid, tannins) that prevented successful extraction of DNA or library preparation. Table 1 has the sample sizes (N) for each study species before and after this filtering.

### Microbiota composition analyses

Statistical analyses and data visualization were completed in R unless otherwise stated. To visualize microbiota composition, stacked barplots were constructed in ggplot2 (v.3.3.2) [92]. These plots showed the bacterial phyla, families, and genera with average relative abundances greater than 1% across samples. We also identified the ASVs that were present in >90% of samples across all host species, and the relative abundances of these ASVs were displayed as heatmaps using the R pheatmap package (v.1.0.12) [93].Sequences from the 10 ASVs were blasted against the National Center for Biotechnology Information (NCBI) Nucleotide database [60] to find similar biological sequences from known bacterial taxa. Furthermore, we also identified ASVs that were biased towards particular host species; these were ASVs that were present in >75% of the samples for a particular host species (e.g. giraffes) and absent in 97% of samples from the other host species.

To detect the bacterial taxa strongly associated with particular host families or dietary guilds, we used the R indicspecies package (v.1.7.9) [94], which calculates an indicator value for each bacterial taxon based on its prevalence in a given group and absence in others. A table of bacterial family relative abundances was used as input, and significance was assessed via permutation tests using 999 permutations (α=0.05). Bacterial families with indicator values >0.4 were plotted in ggplot2.

### Microbiota α-diversity statistical analyses

Prior to alpha-diversity analyses, we controlled for the potential influences of sequencing depth by subsampling all samples to 17,000 sequences using the mothur (v.1.42.3) [95] sub.sample command. Four fecal samples did not meet this sequence cutoff criterion, and were excluded from all alpha-diversity analyses. Mothur was used to construct rarefaction curves of ASV richness vs. sequencing depth (Figure S6) and Good’s coverage values averaged 97.78 ± 0.91 across all samples, indicating that sample coverage was high and appropriate for characterizing fecal microbiota profiles. These values are comparable to those typically reported in other mammalian gut microbiota studies [8, 96, 97].

Microbiota alpha-diversity was estimated using Chao1 Richness, Shannon diversity, and Faith’s phylogenetic diversity in R. Chao1 Richness and Shannon indices were calculated using the phyloseq package (v.1.33.0) [98]. To obtain measures of Faith’s Phylogenetic Diversity (PD), we constructed a phylogenetic tree of ASV sequences using phangorn (v. 2.5.5) [99] and calculated PD using the picante package (v.1.8.1) [100]. The effects of predictor variables on each measure of alpha-diversity across all samples were evaluated via linear mixed models (LMMs) using the lme4 package (v.1.1.23) [101], specifying host dietary guild and host family as fixed variables and sample month as a random effect. A similar model was also built for bovid samples, which included host species as a predictor in lieu of host family. The significance of each predictor variable was determined by calculating likelihood ratio Chi-2 test statistics (α=0.05) on the full models using the car package (v.3.0.7) [102]. These tests were followed by TukeyHSD post-hoc tests with Benjamini-Hochberg adjustments to control for multiple comparisons. Boxplots of microbiota alpha-diversity were generated in ggplot2.

To further quantify the influence of host diet on the three metrics of gut microbiota alpha-diversity, we conducted partial Mantel tests in the R vegan package (v.2.5.7) [103]. Specifically, we evaluated whether similarity in gut microbiota alpha-diversity was associated with similarity in dietary C4(%) after taking into account variation due to host phylogenetic relatedness. The 3 matrices used as input were i) a dissimilarity matrix of gut microbiota alpha-diversity, ii) a dissimilarity matrix of host %C4, and iii) a matrix of host divergence times.

### Microbiota β-diversity analyses and testing for phylosymbiosis

In order to determine the relative contributions and amount of variance explained by host predictor variables, permutational multivariate analyses of variance (PERMANOVA) tests based on Bray-Curtis, Jaccard, and Unifrac distance matrices were run using vegan. Bray-Curtis/Jaccard distances were estimated using vegan, whereas weighted and unweighted Unifrac distances were estimated using phyloseq. Bray-Curtis and weighted Unifrac distances take into account the abundances of bacterial taxa while Jaccard and unweighted Unifrac metrics only consider their presence or absence. Both UniFrac metrics utilize information on the phylogenetic diversity of bacterial members when calculating microbiota similarity. PERMANOVA model #1 included sample month, host dietary guild, host family, and host species as predictors (in this order). Model 2 was identical, except it was restricted to the Bovid dataset and excluded the host family term. Model 3 quantified the effect of the host species predictor while controlling for dietary guild (strata=dietary guild). Microbiota similarity and groupings across samples were visualized via Principal Coordinates Analysis (PCoA) plots.

To test for phylosymbiosis, i.e. the congruence between host phylogenetic relatedness and gut microbiota similarity, mean divergence times (mya) were calculated between every pair of host species in R. First, we retrieved 1000 phylogenetic trees that included all species of Artiodactyla and African elephants (*Loxodonta Africana*) from Upham’s et al. (2019) Mammalian supertree [104]. The trees were randomly sampled from the posterior distribution of Upham’s supertrees (Mammals birth-death tip-dated DNA-only trees) using the VertLife online resource (http://vertlife.org/). Each tree was pruned to include only the species in this study, and branch lengths (i.e. divergence times between each pair of host species) were extracted using the R ape package (v.5.4.1) [105]. All 1000 trees showed the same phylogenetic relationships among the study species and matrices of mean divergence times were estimated from those trees. To determine the strength of the phylosymbiosis signal, we conducted mantel tests on gut microbiota dissimilarity matrices and matrices of host divergence times, setting 999 permutations and Spearman correlations. We also ran partial Mantel tests to evaluate the strength of the phylosymbiosis signal while taking into account gut microbiota variation attributable to diet. The 3 matrices used as input were i) a gut microbiota dissimilarity matrix, ii) a matrix of host divergence times, and iii) a dissimilarity matrix of host %C4 (Table S4).

We visualized the phylosymbiosis findings by plotting gut microbiota similarity (0-1) against host phylogenetic divergence time (mya) in ggplot2. We also constructed a consensus phylogeny of our host species and compared it against a dendrogram of gut microbiota dissimilarity, which was calculated using hierarchical clustering with the R stats package [89] and plotted using the ape package.

### Comparisons of Masai Mara and Laikipia herbivores

In order to compare the gut microbiotas of Masai Mara (1°22′19″S, 34° 56′17″E) herbivores to the gut microbiotas of their conspecifics in Laikipia (0°17′33″N, 36° 53′55″E) (∼ 290 km from the Masai Mara), we downloaded all publicly available sequences from Kartzinel *et al*. [24], and combined them with the raw 16S rRNA gene sequences from this study (Masai Mara herbivores). These were then processed together in DADA2. A total of eight herbivore species overlapped between the two studies: African buffalo, domestic cattle, common eland, impala, giraffe, warthog, plains zebra, and African elephant. The majority of samples from Kartizinel *et al*. [24] and from our study were collected during the wet seasons in their respective regions (Table S8), although, in general, Laikipia is more arid than the Masai Mara, with only 300-600mm precipitation annually [28, 106, 107]. For a list of all samples (N=305), and their associated meta data, see the *Availability of data and materials* section.

The bioinformatics processing and statistical analyses were performed as described above, with a few exceptions. In DADA2, forward and reverse reads were trimmed to 240bp and 150bp, respectively, to better account for sequence quality. Up to 2 errors were allowed per forward read and up to 4 errors per reverse read. To identify the strongest predictors of gut microbiota structure, we constructed a PERMANOVA model that included sample month, host geographic region, host dietary guild, and host species as variables (in this order). PCoA ordinations and testing for phylosymbiosis were conducted as described earlier. To visualize gut microbiota compositions between Masai Mara and Laikipia herbivores, a heatmap of the 32 most abundant bacterial ASVs was constructed using R pheatmap. To statistically evaluate whether conspecific hosts differed in their abundances of these 32 ASVs, we ran 32 distinct LMMs in R, each specifying an ASV relative abundance as the dependent variable, host region as a fixed effect and host species as a random effect (to restrict analyses to within a host species). p-values were adjusted for multiple comparisons using the Bonferroni-Hochberg method. To extend this analysis to include more ASVs, we conducted Linear discriminant analysis Effect Size (LEfSe) [108] in the Galaxy platform [109] using default parameters. Only ASVs >0.01% average relative abundance across samples were included in the dataframe uploaded to Galaxy. ASVs that were enriched in hosts from one geographic region relative to the other were visualized via diverging dot plots in R with the ggplot2 package.

## List of abbreviations used in this manuscript

(ASV): Amplicon Sequence Variant
(LMM): Linear Mixed Model
(PERMANOVA): Permutational Multivariate Analysis of Variance
(PCoA): Principal Coordinates Analysis
(LEfSe): Linear discriminant analysis Effect Size
(NCBI): National Center for Biotechnology Information
(SCFAs): Short Chain Fatty Acids
(UIC): University of Chicago at Illinois
(DADA2): Divisive Amplicon Denoising Algorithm
(PD): Faith’s Phylogenetic Diversity
(mya): Millions of years
(IACUC): Institutional Animal Care and Use Committee

## DECLARATIONS

### Ethical Approval and Consent to participate

Our research and procedures were most recently approved on April 16, 2019 (IACUC approval no. PROTO201900126) and comply with the ethical standards of Michigan State University and Kenya.

## Consent for publication

Not applicable

### Availability of data and materials

The 16S rRNA gene sequence data from this study were deposited in NCBI’s Sequence Read Archive under BioProject PRJNA656793 and accession numbers SAMN15803511-SAMN15803691. Sample metadata, data output by DADA2 (ASV table & ASV taxonomic classifications), data obtained from LEfSe, and R scripts for all analyses and figures included in this manuscript are available on Github (https://github.com/rojascon/Rojas_et_al_2020_African_herbivores_gut_microbiome).

## Supplementary Materials

Supplementary tables and figures can be found online at bioRxiv, alongside this manuscript submission.

### Funding

This work was supported by funds from NSF grants OISE1853934, IOS 1755089, and OIA0939454 to K.E.H. and colleagues, the latter via the BEACON Center for the Study of Evolution in Action. The corresponding author (C.A.R.) was also supported by a Graduate Research Fellowship from NSF, a Predoctoral Fellowship from the Ford Foundation, and a summer fellowship awarded by the MSU Ecology, Evolution, and Behavior program.

## Author Contributions

K.R.T., K.E.H., and C.A.R. designed the study, C.A.R. and Mara Hyena Project field assistants collected the samples. C.A.R., S.R.B., and K.R.T. analyzed the data. C.A.R., K.R.T, and K.E.H. wrote the manuscript and all authors approved the final version of the manuscript.

## Acknowledgements

We thank the field assistants and graduate students from the Mara Hyena Project for assisting with fecal sample collection in the field. We also thank the Kenya Wildlife Service, the Kenyan National Commission on Science, Technology and Innovation, the Kenyan National Environmental Management Authority, the Narok County Government, the Naboisho Conservancy, the Mara Conservancy, and Brian Heath for allowing us to conduct this research.

## Author’s information

Not applicable

## Additional files

**Additional file 1. Tables S1-S8. Table S1-S3:**Multiple-comparison testing of gut microbiota alpha-diversity among host families, species, and dietary guilds. **Table S4:** Previously published dietary %C4 data for the surveyed herbivores. **Table S5:** PERMANOVA tests that included sample month, geographic region, host dietary guild and host species for the combined Masai Mara and Laikipia dataset. **Table S6:** Linear models testing whether conspecific hosts differ in the relative abundances of 32 dominant ASVs. **Table S7:** Proportion of ASVs that were differentially enriched in herbivores from one geographic region compared to the other according to LEfSe. **Table S8:** Sample sizes for each month for the combined Masai Mara and Laikipia dataset (.xlxs 29 KB).

**Additional file 2. Figures S1-S6. Figure S1-S2:** Stacked bar plots showing relative abundances of top bacterial phyla and genera. **Figure S3:** Heatmap of the relative abundances of 10 widely-shared bacterial ASVs. **Figure S4:** PCoA ordinations showing sample clustering within a dietary guild (grazers, browsers, mixed-feeders). **Figure S5:** Stacked barplot of the proportion of samples from each host species that were collected each month (for the Masai Mara dataset). **Figure S6:** Rarefaction curves of ASV richness for the study samples (.pdf 1.9 MB).

## References

1. Jovel J, Dieleman LA, Kao D, Mason AL, Wine E. The Human Gut Microbiome in Health and Disease. Metagenomics. 2018;:197–213. doi:10.1016/B978-0-08-102268-9.00010-0.

2. Ungerfeld E, Leigh M, Forster R, Barboza P. Influence of Season and Diet on Fiber Digestion and Bacterial Community Structure in the Rumen of Muskoxen (Ovibos moschatus). Microorganisms. 2018;6:89. doi:10.3390/microorganisms6030089.

3. Greene LK, McKenney EA, O’Connell TM, Drea CM. The critical role of dietary foliage in maintaining the gut microbiome and metabolome of folivorous sifakas. Sci Rep. 2018;8:1–13.

4. den Besten G, van Eunen K, Groen A, Venema K, Reijngoud D, Bakker B. The role of short-chain fatty acids in the between diet, gut microbiota, and host energy metabolism. J Lipid Res. 2013;54:2325–40. doi:10.1194/jlr.R036012.

5. Park W. Gut microbiomes and their metabolites shape human and animal health. J Microbiol. 2018;56:151–3. doi:10.1007/s12275-018-0577-8.

6. Münger E, Montiel-Castro AJ, Langhans W, Pacheco-López G. Reciprocal Interactions Between Gut Microbiota and Host Social Behavior. Front Integr Neurosci. 2018;12:1–14. doi:10.3389/fnint.2018.00021.

7. Pennisi E. Gut bacteria linked to mental well-being and depression. Science (80-). 2019;363:569–569. doi:10.1126/science.363.6427.569.

8. Li J, Zhan S, Liu X, Lin Q, Jiang J, Li X. Divergence of Fecal Microbiota and Their Associations With Host Phylogeny in Cervinae. Front Microbiol. 2018;9:1–11. doi:10.3389/fmicb.2018.01823.

9. Amato KR, Sanders J, Song SJ, Nute M, Metcalf JL, Thompson LR, et al. Evolutionary trends in host physiology outweigh dietary niche in structuring primate gut microbiomes. ISME J. 2019;13:576–587. doi:10.1038/s41396-018-0175-0.

10. Nishida AH, Ochman H. Rates of gut microbiome divergence in mammals. Mol Ecol. 2018;27:1884–97.

11. Youngblut ND, Reischer GH, Walters W, Schuster N, Walzer C, Stalder G, et al. Host diet and evolutionary history explain different aspects of gut microbiome diversity among vertebrate clades. Nat Commun. 2019;10:1–15. doi:10.1038/s41467-019-10191-3.

12. Moeller AH, Suzuki TA, Lin D, Lacey EA, Wasser SK, Nachman MW. Dispersal limitation promotes the diversification of the mammalian gut microbiota. PNAS. 2017;114:13768–73. doi:10.1073/pnas.1700122114.

13. Kohl KD. Ecological and evolutionary mechanisms underlying patterns of phylosymbiosis in host-associated microbial communities. Philos Trans R Soc B Biol Sci. 2020;375.

14. Lim SJ, Bordenstein SR. An introduction to phylosymbiosis. Proc R Soc B Biol Sci. 2020;287.

15. Brooks AW, Kohl KD, Brucker RM, Van Opstal EJ, Bordenstein SR. Phylosymbiosis: Relationships and Functional Effects of Microbial Communities across Host Evolutionary History. PLoS Biol. 2016;14:1–19. doi:10.1371/journal.pbio.2000225.

16. Watson SE, Hauffe HC, Bull M, McKinney MA, Atwood TA, Perkins SE. Global change-driven use of onshore habitat impacts polar bear faecal microbiota. ISME J. 2019. doi:10.1038/s41396-019-0480-2.

17. Angelakis E, Yasir M, Bachar D, Azhar EI, Lagier JC, Bibi F, et al. Gut microbiome and dietary patterns in different Saudi populations and monkeys. Sci Rep. 2016;6:1–9. doi:10.1038/srep32191.

18. Goertz S, Menezes AB De, Birtles RJ, Id JF, Lowe E, Maccoll ADC, et al. Geographical location influences the composition of the gut microbiota in wild house mice (Mus musculus domesticus) at a fine spatial scale. PLoS One. 2019;14:1–16.

19. Orkin JD, Campos FA, Guadamuz A, Melin AD, Myers MS, Hernandez SEC. Seasonality of the gut microbiota of free-ranging white-faced capuchins in a tropical dry forest. ISME J. 2019;13:183–196. doi:10.1038/s41396-018-0256-0.

20. Hillerislambers J, Adler PB, Harpole WS, Levine JM, Mayfield MM. Rethinking Community Assembly through the Lens of Coexistence Theory. Annu Rev Ecol Evol Syst. 2012;43:227–48. doi:10.1146/annurev-ecolsys-110411-160411.

21. Smith B, Wilson JB. Community convergence: Ecological and evolutionary. Folia Geobot. 2002;37:171–83.

22. Knowles SCL, Eccles RM, Baltrūnaitė L. Species identity dominates over environment in shaping the microbiota of small mammals. Ecol Lett. 2019;22:826–37.

23. Kohl KD, Varner J, Wilkening JL, Dearing MD. Gut microbial communities of American pikas (Ochotona princeps): Evidence for phylosymbiosis and adaptations to novel diets. J Anim Ecol. 2018;87:323–30.

24. Kartzinel TR, Hsing JC, Musili PM, Brown BRP, Pringle RM. Covariation of diet and gut microbiome in African megafauna. PNAS. 2019;116:1–6.

25. Ottichilo WK, De Leeuw J, Skidmore AK, Prins HHT, Said MY. Population trends of large non-migratory wild herbivores and livestock in the Masai Mara ecosystem, Kenya, between 1977 and 1997. Afr J Ecol. 2000;38:202–16.

26. Goheen JR, Augustine DJ, Veblen KE, Kimuyu DM, Palmer TM, Porensky LM, et al. Conservation lessons from large-mammal manipulations in East African savannas: The KLEE, UHURU, and GLADE experiments. Ann N Y Acad Sci. 2018;1429:31–49.

27. Veldhuis MP, Ritchie ME, Ogutu JO, Morrison TA, Beale CM, Estes AB, et al. Cross-boundary human impacts compromise the Serengeti-Mara ecosystem. Science (80-). 2019;363:1424–8. doi:10.1126/science.aav0564.

28. Augustine DJ. Response of native ungulates to drought in semi-arid Kenyan rangeland. Afr J Ecol. 2010;48:1009–20.

29. Broten MD, Said M. Population Trends of Ungulates in and around Kenya’s Masai Mara Reserve. In: Serengeti II: Dynamics, Management, and Conservation of an Ecosystem. 1995. p. 169–93.

30. Cerling TE, Harris JM, Passey BH. Diets of East African Bovidae based on stable isotope analysis. J Mammal. 2003;84:456–70.

31. Kartzinel TR, Chen PA, Coverdale TC, Erickson DL, Kress WJ, Kuzmina ML, et al. DNA metabarcoding illuminates dietary niche partitioning by African large herbivores. PNAS. 2015;112:8019–24.

32. Codron D, Codron J, Lee-Thorp JA, Sponheimer M, De Ruiter D, Sealy J, et al. Diets of savanna ungulates from stable carbon isotope composition of faeces. J Zool. 2007;273:21–9.

33. Codron J, Lee-Thorp JA, Sponheimer M, Codron D, Grant R, De Ruiter D. Elephant (Loxodonta Africana) diets in Kruger National Park, South Africa: Spatial and landscape differences. J Mammal. 2006;87:27–34.

34. Ley RE, Hamady M, Lozupone C, Turnbaugh PJ, Ramey RR, Bircher JS, et al. Evolution of Mammals and Their Gut Microbes. Science (80-). 2008;320:1647.

35. McKenzie VJ, Song SJ, Delsuc F, Prest TL, Oliverio AM, Korpita TM, et al. The effects of captivity on the mammalian gut microbiome. Integr Comp Biol. 2017;57:690– 704.

36. Phillips CD, Phelan G, Dowd SE, McDonough MM, Ferguson AW, Delton Hanson J, et al. Microbiome analysis among bats describes influences of host phylogeny, life history, physiology and geography. Mol Ecol. 2012;21:2617–27.

37. Groussin M, Mazel F, Sanders JG, Smillie CS, Lavergne S, Thuiller W, et al. Unraveling the processes shaping mammalian gut microbiomes over evolutionary time. Nat Commun. 2017;8:14319. doi:10.1038/ncomms14319.

38. Groussin M, Mazel F, Alm EJ. Co-evolution and Co-speciation of Host-Gut Bacteria Systems. Cell Host Microbe. 2020;28:12–22. doi:10.1016/j.chom.2020.06.013.

39. Godon JJ, Arulazhagan P, Steyer JP, Hamelin J. Vertebrate bacterial gut diversity: Size also matters. BMC Ecol. 2016;16:1–9.

40. Liu P, Cheng A, Huang S, Lu H, Oshida T. Body-size Scaling is Related to Gut Microbial Diversity, Metabolism and Dietary Niche of Arboreal Folivorous Flying Squirrels. Sci Rep. 2020;:1–12. doi:10.1038/s41598-020-64801-y.

41. Budd K, Gunn JC, Eggert LS, Finch T, Klymus K, Sitati N. Effects of diet, habitat, and phylogeny on the fecal microbiome of wild African savanna (Loxodonta africana) and forest elephants (L. cyclotis). Ecol Evol. 2020;00:1–14.

42. Metcalf JL, Song SJ, Morton JT, Weiss S, Seguin-Orlando A, Joly F, et al. Evaluating the impact of domestication and captivity on the horse gut microbiome. Sci Rep. 2017;7.

43. Illius AW, Gordon IJ. The functional significance of the browser-grazer dichotomy in African ruminants. Oecologia. 1994;98:167–75.

44. Miller EA, Livermore J, Alberts SC, Tung J, Archie EA. Ovarian cycling and reproductive state shape the vaginal microbiota in wild baboons. Microbiome. 2017;5:1– 14.

45. Moeller AHA, Foerster S, Wilson ML, Pusey AE, Hahn BH, Ochman H. Social behavior shapes the chimpanzee pan-microbiome. Sci Adv. 2016;2:e1500997– e1500997. doi:10.1126/sciadv.1500997.

46. Johnson KVA. Gut microbiome composition and diversity are related to human personality traits. Hum Microbiome J. 2020;15 December 2019:100069. doi:10.1016/j.humic.2019.100069.

47. Shipley L. Grazers and browsers: how digestive morphology affects diet selection. Grazing Behav Livest Wildl. 1999;:20–7. http://www.cnr.uidaho.edu/range456/readings/shipley.pdf.

48. Venter JA, Vermeulen MM, Brooke CF. Feeding Ecology of Large Browsing and Grazing Herbivores. In: Gordon IJ, Prins HHT, editors. The Ecology of Browsing and Grazing II. Cham: Springer International Publishing; 2019. p. 127–53. doi:10.1007/978-3-030-25865-8_5.

49. Hofmann RR. Evolutionary steps of ecophysiological adaptation and diversification of ruminants: a comparative view of their digestive system. Oecologia. 1989;78:443–57. doi:10.1007/BF00378733.

50. Spencer LM. Morphological Correlates of Dietary Resource Partitioning in the African Bovidae. J Mammal. 1995;76:448–71. doi:10.2307/1382355.

51. Henderson G, Cox F, Ganesh S, Jonker A, Young W, Janssen PH, et al. Rumen microbial community composition varies with diet and host, but a core microbiome is found across a wide geographical range. Sci Rep. 2015;5:14567.

52. Edwards JE, Shetty SA, Berg P Van Den, Burden F, Doorn DA Van, Pellikaan WF, et al. Multi-kingdom characterization of the core equine fecal microbiota based on multiple equine (sub) species. Anim Microbiome. 2020;2:1–16.

53. Ilmberger N, Güllert S, Dannenberg J, Rabausch U, Torres J, Wemheuer B, et al. A comparative metagenome survey of the fecal microbiota of a breast-and a plant-fed asian elephant reveals an unexpectedly high diversity of glycoside hydrolase family enzymes. PLoS One. 2014;9:1–12.

54. Roggenbuck M, Sauer C, Poulsen M, Bertelsen MF, Sørensen SJ. The giraffe (Giraffa camelopardalis) rumen microbiome. FEMS Microbiol Ecol. 2014;90:237–46.

55. Tan SC, Chong CW, Yap IKS, Thong KL, Teh CSJ. Comparative assessment of faecal microbial composition and metabonome of swine, farmers and human control. Sci Rep. 2020;10:1–13.

56. Zeng B, Zhang S, Xu H, Kong F, Yu X, Wang P, et al. Gut microbiota of Tibetans and Tibetan pigs varies between high and low altitude environments. Microbiol Res. 2020;235 November 2019:126447. doi:10.1016/j.micres.2020.126447.

57. Biddle A, Stewart L, Blanchard J, Leschine S. Untangling the genetic basis of fibrolytic specialization by Lachnospiraceae and Ruminococcaceae in diverse gut communities. Diversity. 2013;5.

58. Flint HJ, Scott KP, Duncan SH, Louis P, Forano E. Microbial degradation of complex carbohydrates in the gut. Gut Microbes. 2012; August:289–306.

59. Kohl KD, Denise Dearing M. The woodrat gut microbiota as an experimental system for understanding microbial metabolism of dietary toxins. Front Microbiol. 2016;7:1–9.

60. Coordinators NR. Database resources of the National Center for Biotechnology Information. Nucleic Acids Res. 2018;46. doi:10.1093/nar/gkx1095.

61. Scheelings TF, Moore RJ, Van TTH, Klaassen M, Reina RD. Microbial symbiosis and coevolution of an entire clade of ancient vertebrates: the gut microbiota of sea turtles and its relationship to their phylogenetic history. Anim Microbiome. 2020;2:2–12.

62. Tinker KA, Ottesen EA. Phylosymbiosis across deeply diverging lineages in omnivorous cockroaches. Appl Environ Microbiol. 2020;86:e02513–19. doi:10.1128/AEM.02513-19.

63. Schmidt E, Mykytczuk N, Schulte-Hostedde AI. Effects of the captive and wild environment on diversity of the gut microbiome of deer mice (Peromyscus maniculatus). ISME J. 2019;13:1293–305. doi:10.1038/s41396-019-0345-8.

64. Perofsky AC, Lewis RJ, Meyers LA. Terrestriality and bacterial transfer: a comparative study of gut microbiomes in sympatric Malagasy mammals. ISME J. 2019;13:50–63. doi:10.1038/s41396-018-0251-5.

65. Greene LK, Clayton JB, Rothman RS, Semel BP, Semel MA, Gillespie TR, et al. Local habitat, not phylogenetic relatedness, predicts gut microbiota better within folivorous than frugivorous lemur lineages. Biol Lett. 2019;15:5–11.

66. Gogarten JF, Davies TJ, Benjamino J, Gogarten JP, Graf J, Mielke A, et al. Factors influencing bacterial microbiome composition in a wild non-human primate community in Taï National Park, Côte d’Ivoire. ISME J. 2018;12:2559–74. doi:10.1038/s41396-018-0166-1.

67. Amato KR, Mallott EK, Mcdonald D, Dominy NJ, Goldberg T, Lambert JE, et al. Convergence of human and Old World monkey gut microbiomes demonstrates the importance of human ecology over phylogeny. Genome Biol. 2019;20:201.

68. Gerardo NM, Hoang KL, Stoy KS. Evolution of animal Immunity in the light of beneficial symbioses. Philos Trans R Soc B2. 2020;375:20190601.

69. Popkes M, Valenzano DR. Microbiota-host interactions shape ageing dynamics. Philos Trans R Soc B2. 2020;375:20190596.

70. Grieneisen LE, Charpentier MJE, Alberts SC, Blekhman R, Bradburd G, Tung J, et al. Genes, geology and germs: Gut microbiota across a primate hybrid zone are explained by site soil properties, not host species. Proc R Soc B Biol Sci. 2019;286.

71. Muegge BD, Kuczynski J, Kinghts D, Clemente JC, Fontana L, Henrissat B, et al. Diet drives Convergence in Gut Microbiome Functions Across Mammalian Phylogeny and Within Humans. Science (80-). 2017;332:970–4.

72. Clemens ET, Maloiy GMO. Digestive physiology of East African wild ruminants. Comp Biochem Physiol -- Part A Physiol. 1983;76:319–33.

73. Pérez-Barbería JF, Gordon IJ, Illius AW. Phylogenetic analysis of stomach adaptation in digestive strategies in African ruminants. Oecologia. 2001;129:498–508.

74. Shorrocks B. The Biology of African Savannas. New York: Oxford University Press; 2007.

75. Wright ADG, Auckland CH, Lynn DH. Molecular diversity of methanogens in feedlot cattle from Ontario and Prince Edward Island, Canada. Appl Environ Microbiol. 2007;73:4206–10. doi:10.1128/AEM.00103-07.

76. Daniel H, Gholami AM, Berry D, Desmarchelier C, Hahne H, Loh G, et al. High-fat diet alters gut microbiota physiology in mice. ISME J. 2014;8:295–308. doi:10.1038/ismej.2013.155.

77. Liu J, Pu YY, Xie Q, Wang JK, Liu JX. Pectin Induces an In Vitro Rumen Microbial Population Shift Attributed to the Pectinolytic Treponema Group. Curr Microbiol. 2014;70:67–74. doi:10.1007/s00284-014-0672-y.

78. Xie X, Yang C, Guan LL, Wang J, Xue M, Liu JX. Persistence of Cellulolytic Bacteria Fibrobacter and Treponema After Short-Term Corn Stover-Based Dietary Intervention Reveals the Potential to Improve Rumen Fibrolytic Function. Front Microbiol. 2018;9 JUN:1363. doi:10.3389/fmicb.2018.01363.

79. Kennedy K, Hall MW, Lynch MDJ, Moreno-Hagelsieb G, Neufeld JD. Evaluating bias of Illumina-based bacterial 16S rRNA gene profiles. Appl Environ Microbiol. 2014;80:5717–22.

80. De La Cuesta-Zuluaga J, Escobar JS. Considerations For Optimizing Microbiome Analysis Using a Marker Gene. Front Nutr. 2016;3:26. doi:10.3389/fnut.2016.00026.

81. Mallott EK, Malhi RS, Amato KR. Assessing the comparability of different DNA extraction and amplification methods in gut microbial community profiling. Access Microbiol. 2019.

82. Green DS, Zipkin EF, Incorvaia DC, Holekamp KE. Long-term ecological changes influence herbivore diversity and abundance inside a protected area in the Mara-Serengeti ecosystem. Glob Ecol Conserv. 2019;20:e00697. doi:10.1016/j.gecco.2019.e00697.

83. Kingdon J. East African Mammals: Volume I. Chicago: The University of Chicago Press; 1984.

84. Kingdon J. East African Mammals: Volume IIIC. Chicago: The University of Chicago Press; 1982.

85. Kingdon J. East African Mammals: Volume IIID. Chicago: The University of Chicago Press; 1982.

86. Kingdon J. East African Mammals: Volume IIIB. Chicago: The University of Chicago Press; 1979.

87. Caporaso JG, Lauber CL, Walters WA, Berg-Lyons D, Huntley J, Fierer N, et al. Ultra-high-throughput microbial community analysis on the Illumina HiSeq and MiSeq platforms. ISME J. 2012;6:1621–4. doi:10.1038/ismej.2012.8.

88. Kozich JJ, Westcott SL, Baxter NT, Highlander SK, Schloss PD. Development of a Dual-Index Sequencing Strategy and Curation Pipeline for Analyzing Amplicon Sequence Data on the MiSeq Illumina Sequencing Platform. Appl Environ Microbiol. 2013;79:5112–20. doi:10.1128/AEM.01043-13.

89. R Core Team. R: a language and environment for statistical computing. In: R Foundation for Statistical Computing. Vienna, Austria; 2019. URLhttps://www.gbif.org/tool/81287/r-a-language-and-environment-for-statistical-computing. Accessed 28 Aug 2019.

90. Callahan BJ, McMurdie PJ, Rosen MJ, Han AW, Johnson AJA, Holmes SP. DADA2: High-resolution sample inference from Illumina amplicon data. Nat Methods. 2016;13:581–3.

91. Quast C, Pruesse E, Yilmaz P, Gerken J, Schweer T, Yarza P, et al. The SILVA ribosomal RNA gene database project: improved data processing and web-based tools. Nucleic Acids Res. 2013;41:D590–6. doi:10.1093/nar/gks1219.

92. Wickham H. ggplot2: Elegant Graphics for Dat Analysis. New York, NY: Springer-Verlag New York; 2009. doi:10.1007/978-0-387-98141-3.

93. Kolde R. Pheatmap: Pretty Heatmaps. R package version 1.0.12. 2019;:1–8. URLhttps://cran.r-project.org/package=pheatmap.

94. De Cáceres M, Legendre P. Associations between species and groups of sites: Indices and statistical inference. Ecology. 2009;90:3566–74.

95. Schloss PD, Westcott SL, Ryabin T, Hall JR, Hartmann M, Hollister EB, et al. Introducing mothur: open-source, platform-independent, community-supported software for describing and comparing microbial communities. Appl Environ Microbiol. 2009;75:7537–41. doi:10.1128/AEM.01541-09.

96. Jin L, Wu D, Li C, Zhang A, Xiong Y, Wei R, et al. Bamboo nutrients and microbiome affect gut microbiome of giant panda. Symbio. 2020;80:293–304.

97. Bo T-B, Zhang X-Y, Wen J, Deng K, Qin X-W, Wang D-H. The microbiota-gut-brain interaction in regulating host metabolic adaptation to cold in male Brandt’s voles (Lasiopodomys brandtii). ISME J. 2019. doi:10.1038/s41396-019-0492-y.

98. McMurdie PJ, Holmes S. Phyloseq: An R Package for Reproducible Interactive Analysis and Graphics of Microbiome Census Data. PLoS One. 2013;8:1–11.

99. Schliep KP. phangorn: Phylogenetic analysis in R. Bioinformatics. 2011;27:592–3.

100. Kembel SW, Cowan PD, Helmus MR, Cornwell WK, Morlon H, Ackerly DD, et al. Picante: R tools for integrating phylogenies and ecology. Bioinformatics. 2010;26:1463– 4.

101. Bates D, Mächler M, Bolker B, Walker S. Fitting Linear Mixed-Effects Models Using lme4. J Stat Softw. 2015;67:1–48. doi:10.18637/jss.v067.i01.

102. Fox J, Weisberg S, Fox J. An R companion to applied regression. Thousand Oaks, CA: Sage; 2011.

103. Oksanen J, Blanchet F, Friendly M, Kindt R, Legendre P, McGlinn D, et al. vegan: Community Ecology Package. R Packag version 24-6. 2018. URLhttps://cran.r-project.org/web/packages/vegan/index.html. xAccessed 28 Aug 2019.

104. Upham N, Esselstyn J, Jetz W. Inferring the mammal tree: species-level sets of phylogenies for questions in ecology, evolution, and conservation. PLoS Biol. 2019;17:e3000494.

105. Paradis E, Claude J, Strimmer K. APE: Analyses of phylogenetics and evolution in R language. Bioinformatics. 2004;20:289–90.

106. Ochieng EO. Characterizing the spatial distributions of elephants in Mpala, Kenya. 2015.

107. Kartzinel TR, Pringle RM. Multiple dimensions of dietary diversity in large mammalian herbivores. J Anim Ecol. 2020;0:1–15.

108. Segata N, Izard J, Waldron L, Gevers D, Miropolsky L, Garrett WS, et al. Metagenomic biomarker discovery and explanation. Genome Biol. 2011;12:R60. doi:10.1186/gb-2011-12-6-r60.

109. Afgan E, Baker D, Batut B, van den Beek M, Bouvier D, Čech M, et al. The Galaxy platform for accessible, reproducible and collaborative biomedical analyses: 2018 update. Nucleic Acids Res. 2018;46:W537–44. doi:10.1093/nar/gky379.

