## Supplementary Figures for "Host phylogeny and host ecology structure the mammalian gut microbiota at different taxonomic scales"

**Figure S1. Predominant bacterial phyla of African herbivore gut microbiotas.** Stacked bar plots showing the relative percentage of 16S rRNA gene sequences assigned to each bacterial phylum across samples. Samples are grouped by host species, and each color represents a bacterial phylum.

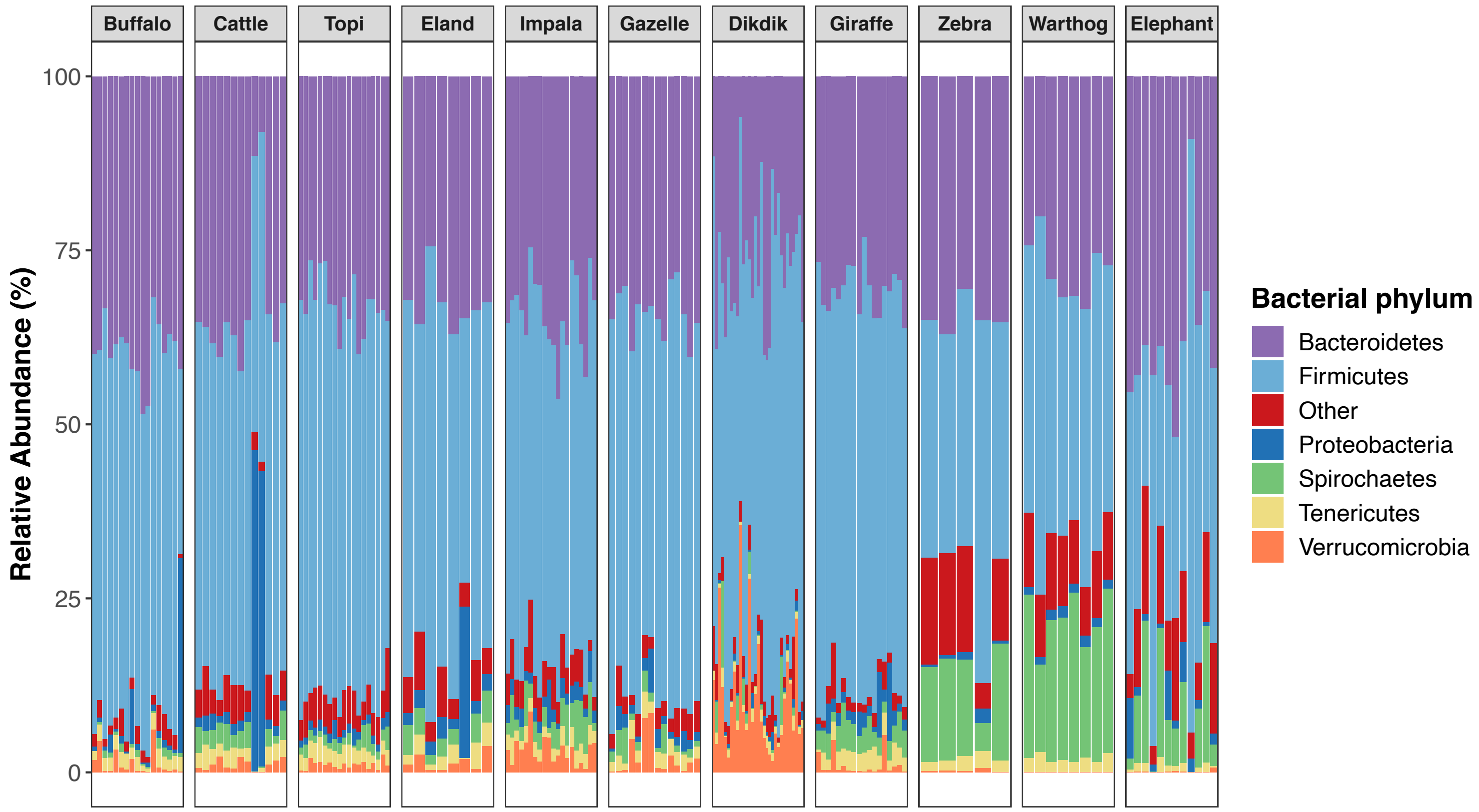

**Figure S2. Predominant bacterial genera of African herbivore gut microbiotas.** Stacked bar plots showing the relative frequency of 16S rRNA gene sequences assigned to each bacterial genus across samples. Samples are grouped by host species, and each color represents a bacterial genus.

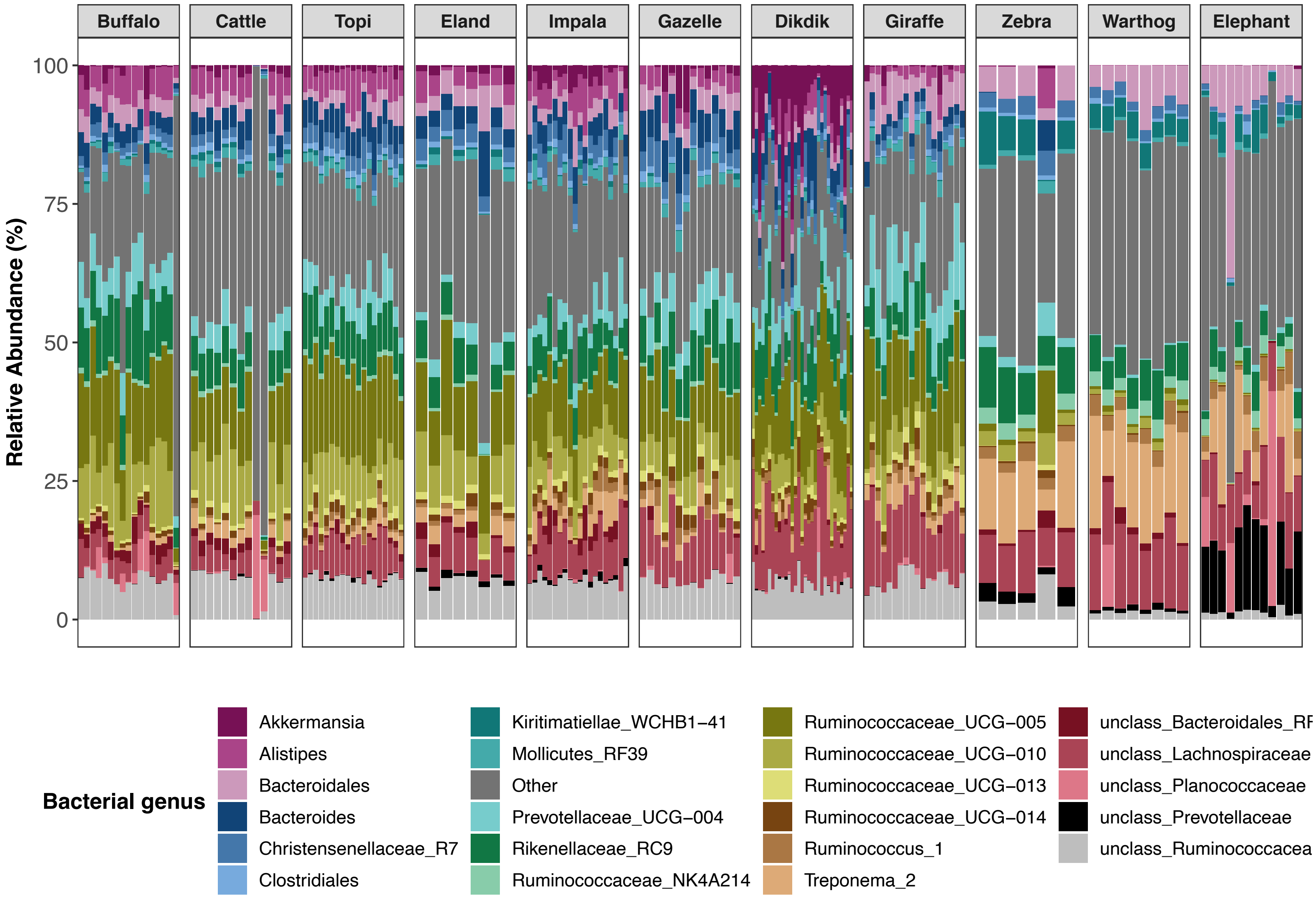

**Figure S3. Relative abundance of 10 ASVs widely shared across host species.** Heatmap showing the relative abundances (proportions) of 10 ASVs that were present in over 90% of the samples included in this study. Samples are grouped and color-coded by host species. Darker colors in the heatmap indicate higher relative abundances.

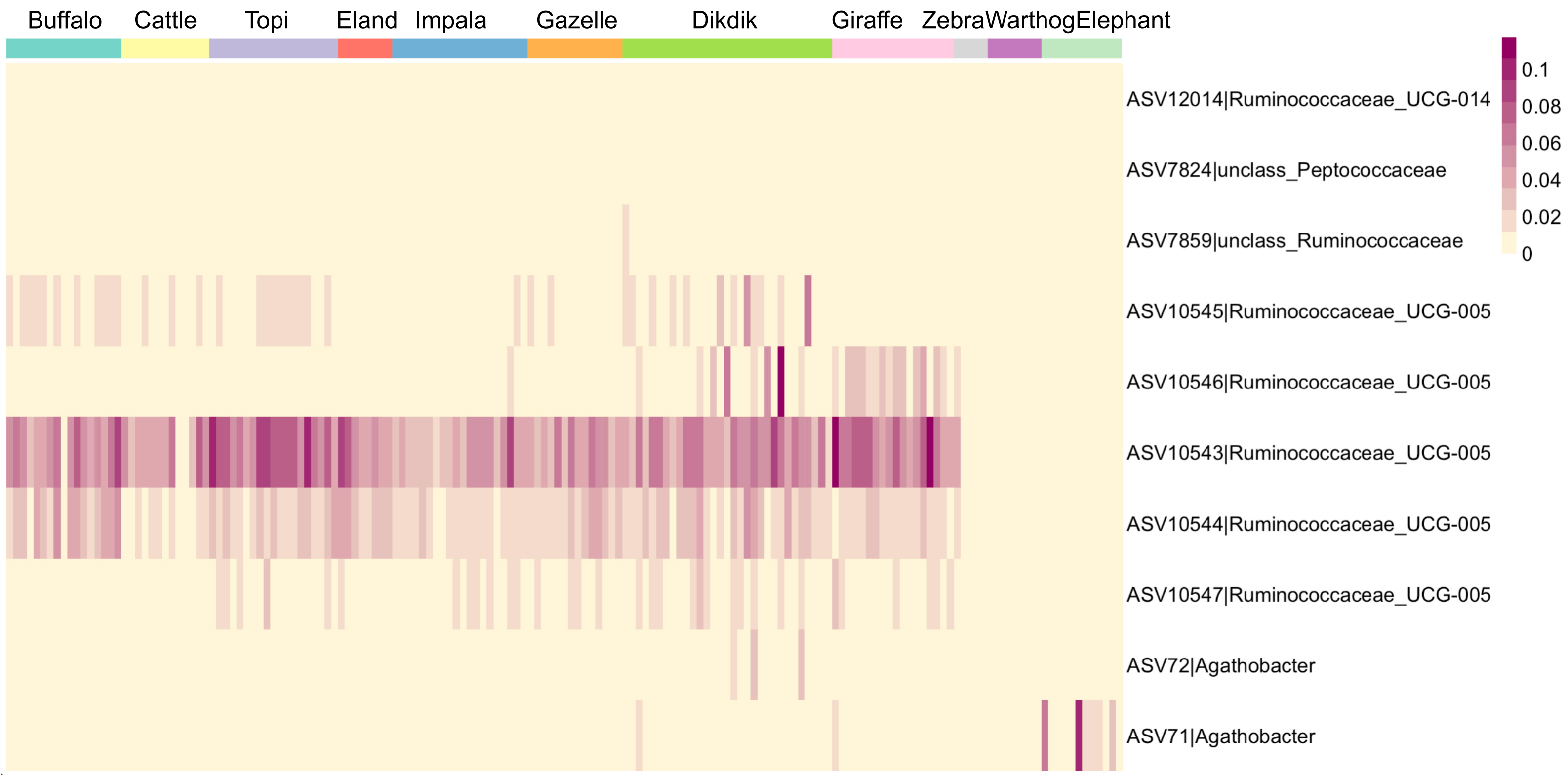


**Figure S4. Within dietary guilds, the gut microbiota is highly species-specific.** PCoA plots constructed from Bray-Curtis dissimilarity matrices. Each point represents a sample and is color-coded by host species. Samples are grouped by dietary guild: grazers (left), browsers (middle), and mixed-feeders (right). Closeness of points indicates high community similarity. The % of variance accounted for by each principal-coordinate axis is shown in the axis labels. PERMANOVA analyses show that host species accounts for 40% of observed variation.

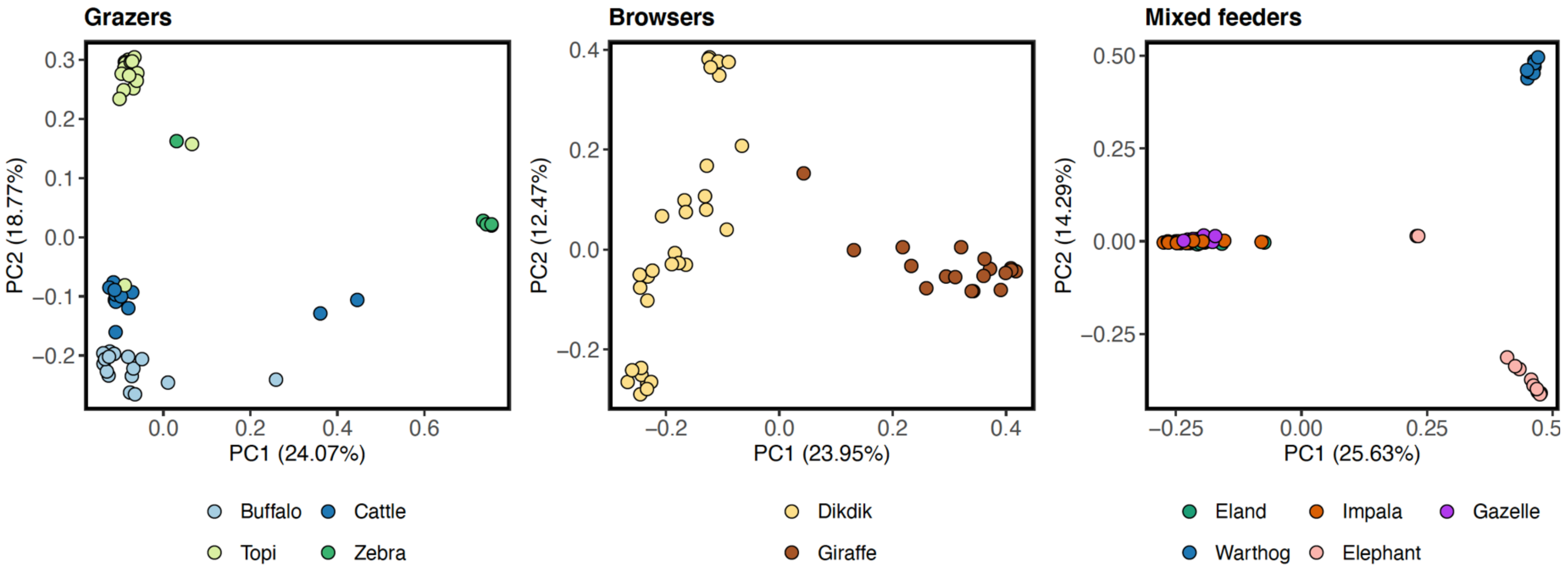

**Figure S5. Sample distribution by Month for Masai Mara herbivores.** The total number of samples collected for each host species at each sampling month is shown.

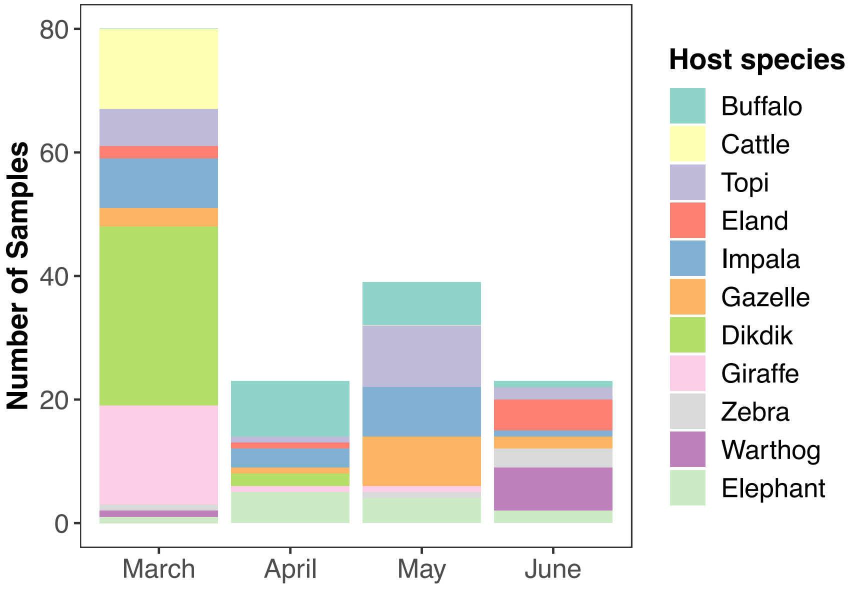


**Figure S6. Rarefaction curves of gut microbiota ASV richness.** Plotted are the number of ASVs (ASV Richness) that are recovered with an increasing number of sequences, after subsampling to 17,000 sequences/sample. Each curve represents a unique sample and is color-coded by host species. The y and x-axis are scaled equally to show what the curves would look if with each read came a new ASV. The curves all plateau.

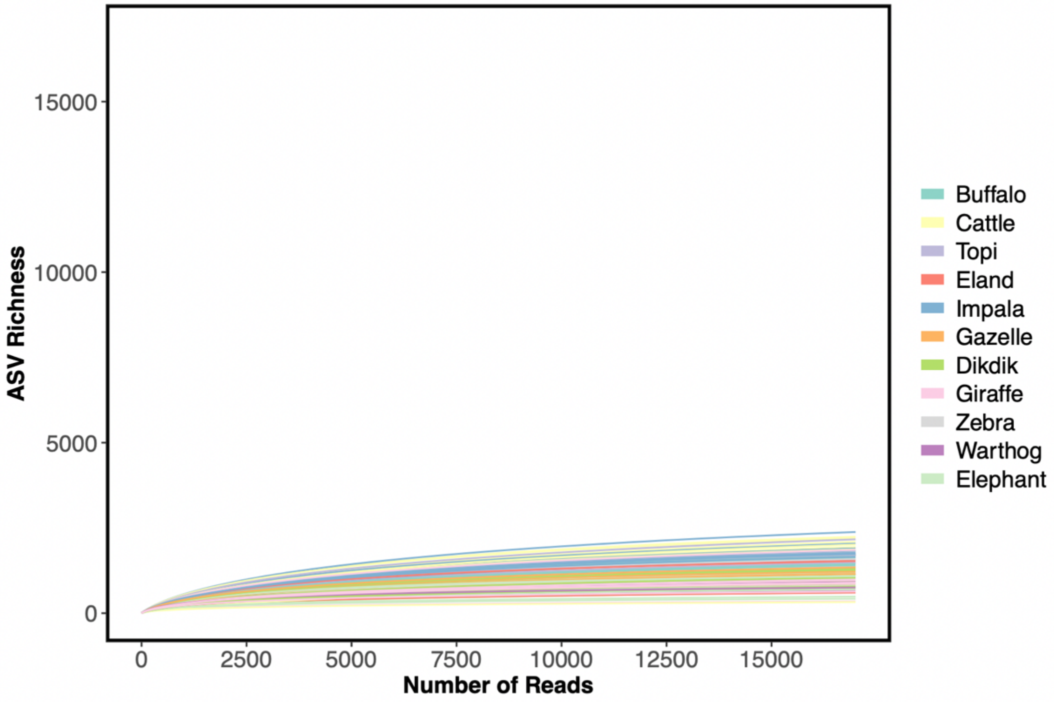
